# Drug-target binding quantitatively predicts optimal antibiotic dose levels

**DOI:** 10.1101/369975

**Authors:** Fabrizio Clarelli, Adam Palmer, Bhupender Singh, Merete Storflor, Silje Lauksund, Ted Cohen, Sören Abel, Pia Abel zur Wiesch

## Abstract

Combatting antibiotic resistance will require both new antibiotics and strategies to preserve the effectiveness of existing drugs. Both approaches would benefit from predicting optimal dosing of antibiotics based on drug-target binding parameters that can be measured early in drug development and that can change when bacteria become resistant. This would avoid the currently frequently employed trial-and-error approaches and might reduce the number of antibiotic candidates that fail late in drug development.

Here, we describe a computational model (COMBAT-COmputational Model of Bacterial Antibiotic Target-binding) that leverages accessible biochemical parameters to quantitatively predict antibiotic dose-response relationships. We validate our model with MICs of a range of quinolone antibiotics in clinical isolates demonstrating that antibiotic efficacy can be predicted from drug-target binding (R^2^ > 0.9). To further challenge our approach, we do not only predict antibiotic efficacy from biochemical parameters, but also do the reverse: estimate the magnitude of changes in drug-target binding based on antibiotic dose-response curves. We experimentally demonstrate that changes in drug-target binding can be predicted from antibiotic dose-response curves with 92-94 % accuracy by exposing bacteria overexpressing target molecules to ciprofloxacin. To test the generality of COMBAT, we apply it to a different antibiotic class, the beta-lactam ampicillin, and can again predict binding parameters from dose-response curves with 90 % accuracy. We then apply COMBAT to predict antibiotic concentrations that can select for resistance due to novel resistance mutations.

Our goal here is dual: First, we address a fundamental biological question and demonstrate that drug-target binding determines bacterial response to antibiotics, although antibiotic action involves many additional effects downstream of drug-target binding. Second, we create a tool that can help accelerate drug development by predicting optimal dosing and preserve the efficacy of existing antibiotics by predicting optimal treatment for possible resistant mutants.

## Introduction

The rise of antibiotic resistance represents an urgent public health threat. In order to effectively combat the spread of antibiotic resistance, we must optimize the use of existing drugs and develop new drugs that are effective against drug-resistant strains. Accordingly, methods to improve antibiotic dose levels to i) maximize efficacy against susceptible strains and ii) minimize resistance evolution play a key role in our defense against antibiotic resistant pathogens.

It is noteworthy that dosing strategies for treatment of susceptible strains (e.g., dosing level[1], dosing frequency[2], and treatment duration[3–5]) have recently been substantially improved, even for antibiotic treatments that have been standard of care for decades. This suggests that there likely remains significant room for optimization in our antibiotic treatment regimens. It also highlights the difficulty in identifying optimal dosing levels for new antibiotics. Indeed, optimizing dosing is one of the biggest challenges in drug development. Typically, time-consuming trial-and-error approaches are used and each failed drug candidate makes this process more expensive[6].

It is even more challenging to optimize dose levels to minimize the emergence of antibiotic resistance, both for existing and novel antibiotics. There remains substantial debate about which dosing strategies best prevent the emergence of resistant mutants during treatment[7–9]. In this context, a useful concept that links antibiotic concentrations with resistance evolution is the resistance selection window (mutant selection window) that ranges from the lowest concentration at which the resistant strain grows faster than the wild-type, usually well below the wild-type minimum inhibitory concentration (MIC), to the MIC of the resistant strain[10–12]. Antibiotic concentrations above the resistance selection window safeguard against *de novo* resistance emergence. Antibiotic concentrations below the resistance selection window do not kill the susceptible strain, but also do not favor the resistant strain and therefore do not promote emergence of resistance. The latter may be preferable if one cannot dose above the MIC of the resistant strain due to toxicity or solubility limits. To limit resistance emergence, it is therefore important to identify the resistance selection window and optimize dosing accordingly.

Limitations in our knowledge of how antibiotic treatment regimens affect bacterial populations contribute to the need for lengthy and expensive trial-and-error approaches, with the sheer number of possible dosing regimens making it difficult to identify an optimal regimen. We argue that this knowledge gap is a major limitation for the improvement of dosing regimens of existing drugs and a real obstacle for the development of new antibiotics[13, 14].

Pharmacodynamic models that can make predictions of bacterial killing and selection on the basis of drug-target interactions offer new promise to inform rational antibiotic dosing practices[15]. Recently described models that include drug-target binding have been useful in gaining a better qualitative understanding of complicated drug effects, such as post-antibiotic effects, inoculum effects, and bacterial persistence[15–18]. However, to speed the development of new antibiotics or to inform practices which minimize resistance, we require quantitative predictions for antibiotics or resistant bacterial strains that do not exist yet. Models which permit quantitative predictions of changes in drug efficacy as a function of modification of antibiotic molecules (i.e. new drugs) or novel resistance mutations would be invaluable. Such tools would advance our general mechanistic understanding of antibiotic action, could guide dosing trials of new drugs, and suggest better dosing of existing drugs.

In this report, we describe a mechanistic computational modeling framework (COMBAT-COmputational Model of Bacterial Antibiotic Target-binding) that allows us to predict drug effects based solely on accessible biochemical parameters describing drug-target interaction. These parameters can be determined early in drug development. We use this framework to investigate how changes in drug target binding, either due to improvements in existing antibiotics or due to resistance mutations in bacteria, affect antibiotic efficacy. We first show that COMBAT accurately predicts bacterial susceptibility as a function of drug-target binding and, conversely, allows inference of these biochemical parameters on the basis of observed patterns of bacterial growth suppression or killing. We then use COMBAT to predict the susceptibility of newly arising resistant variants based on the molecular mechanism of resistance and determine the resistance selection window.

## Results

### Quinolone target affinities correlate with antibiotic efficacy

To investigate how biochemical changes in antibiotic action modifies bacterial susceptibility, we explored how the affinity of antibiotics to their target affects the MIC. We compared the MICs of quinolones, an antibiotic class in which individual antibiotics have a wide range of affinities to their target, gyrase (*K*_*D*_~10^−4^ - 10^−7^ M) but are of similar molecular sizes and have a similar mode of action[19]. This choice allowed us to isolate the effects of differences in drug-target affinity on the MIC.

We obtained binding affinities of quinolones to their gyrase target in *Escherichia coli* from previous studies[20–24]. We then retrieved MIC data for several quinolones from clinical Enterobacteriaceae isolates collected before 1990[25], i.e., before the widespread emergence of quinolone resistance[19]. We assume that quinolone affinities obtained from clinical Enterobacteriaceae isolates collected before the emergence of resistance correspond to those measured in wild-type *E. coli*.

To make qualitative predictions of MICs, we employed a simplified model based on the assumptions that i) drug-target binding occurs much more quickly than bacterial replication, ii) the antibiotic concentration remains constant and iii) that during the 18 hours of an MIC assay, the concentration gradient of the drug inside and outside the cell has equilibrated. Under these assumptions, the MIC can be expressed as

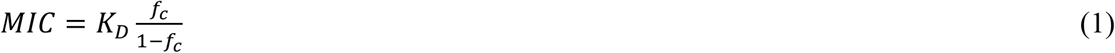

where *K*_*D*_ represents the affinity constant and *f*_*c*_ the fraction of the target bound at the MIC[26]. Accordingly, this model predicts that the MIC is linearly correlated with *K*_*D*_.

Fig. 1 shows the correlations between drug-target affinities and MICs for seven quinolones and clinical isolates of 11 different Enterobacteriaceae species. We observed a significant (*p* < 0.018) linear correlation between MIC and *K*_*D*_ in all species, confirming the qualitative model prediction.

**Fig. 1.**
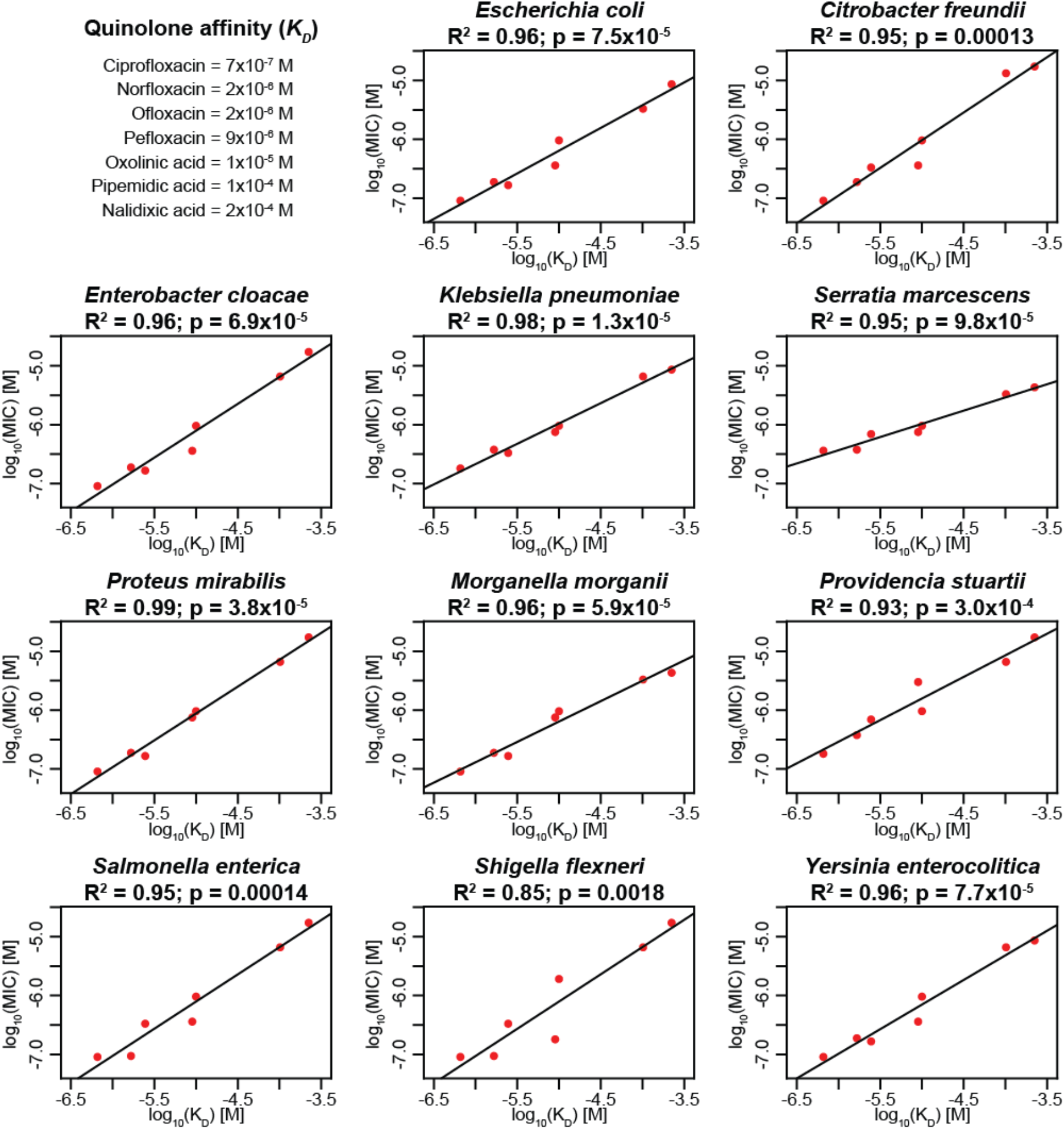
Clinical data confirm linear correlation between MICs and affinities of quinolones to gyrase. We analyzed MIC and drug-target affinity data from 11 Enterobacteriaceae isolates and seven different quinolones. The x-axes show the affinities (*K*_*D*_) as reported in the literature[21, 23–25], and the y-axes show the MICs, both in mol/L. The adjusted R^2^ and p-value of each correlation are given. In cases where there was more than one *K*_*D*_ value reported in the literature, we used the mean for this analysis. The tested MIC values are the median of several clinical isolates described previously[25].

### A quantitative model to predict antibiotic efficacy

While it was encouraging that our model can qualitatively predict MIC changes, our aim was to quantitatively predict antibiotic treatment performance. The simplified model assumes that the binding kinetics are much faster than bacterial replication, which may not be true in all cases. To expand the generalizability of the model, we extended the modeling framework to allow that bacterial replication may occur in a similar time frame as drug-target binding events.

The full model (COMBAT-COmputational Model of Bacterial Antibiotic Target-binding) describes the binding and unbinding of antibiotics to their targets and predicts how such binding dynamics affects bacterial replication and death (Fig. 2a). In previous work linking drug-target binding kinetics with bacterial replication[18], we described a population of bacteria with *θ* target molecules per cell with a system of *θ* + 1 (bacteria with 0, 1, …, *θ* bound target molecules) ordinary differential equations (ODEs). This system increases in complexity with the number of target molecules and makes fitting the model to data computationally too demanding for most settings. To simplify this prior approach, we developed new mathematical models based on partial differential equations (PDEs), where a single equation describes all bacteria simultaneously. The sum of bacteria within all target occupancy states over time can be described by a time kill curve (Fig. 2b), during which the bacterial population is characterized by the distribution of bacterial cells with different levels of target occupancies at each time-step (Fig. 2c). This curve can be visualized as a two-dimensional surface in a three-dimensional coordinate system where the number of bacteria is represented on the z-axis, the percent of bacteria with the fraction of bound target molecules on the x-axis, and time on the y-axis (Fig. 2d).

**Fig. 2.**
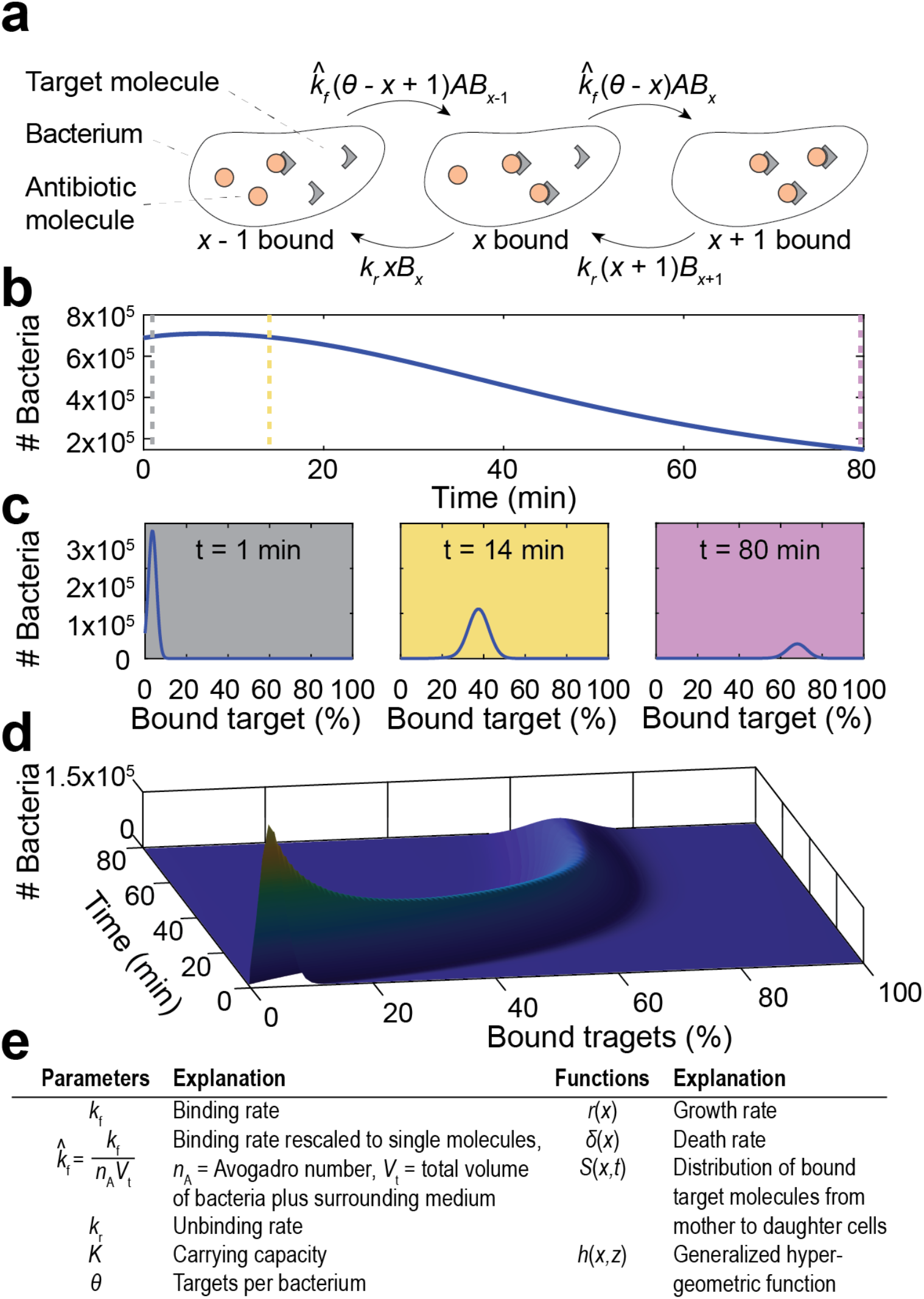
Illustration of modeling approach. **a**, Schematic illustration of binding kinetics (adapted from[52]). The grey triangles depict the drug target molecules, and the orange circles represent antibiotic molecules within bacteria. The arrows indicate individual binding and unbinding events of the antibiotic to its target molecule in the cell. 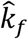 is the adjusted forward reaction rate, *k*_*r*_ is the reverse reaction rate, *A* is the concentration of antibiotics inside the bacterium, *x* is the number of bound targets, *θ* is the number of targets and *B*_*x*_ is the number of bacteria with *x* bound targets. **b**, Modeled sample time-kill curve, in which the sum of bacteria in all binding states (i.e., the entire population of living bacteria) is followed over time after exposure to antibiotics. The vertical dotted lines indicate the time points depicted in (**c**); 1 min (grey), 14 min (yellow), and 80 min (purple). **c**, The percentage of bound antibiotic targets in the bacterial population at indicated time points. **d**, Illustration of how the partial differential equation describes the bacterial population as a surface in a three-dimensional coordinate system, the dimensions of which represent percent bound target (x-axis), time (y-axis), and number of bacteria (z-axis). The three time points shown in (**c**) represent two-dimensional cross-sections at different points of the y-axis. **e**, Overview of used parameters and functions.

Antibiotic action is described by rates of binding (*k*_*f*_) and unbinding (*k*_*r*_) to bacterial target molecules (Fig. 2a, e). The binding of an antibiotic to a target results in the formation of an antibiotic-target molecule complex *x*, where *x* ranges between 0 and *θ*.

COMBAT consists of two mass balance equations: equation 2 describing bacterial numbers as a function of bound targets and time and equation 3 describing antibiotic concentration as a function of time (Methods section).

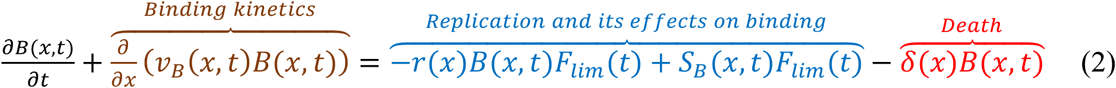

The term for binding kinetics is given in brown, the term for replication in blue and the term for death in red.

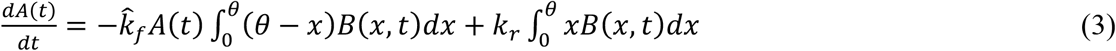

where *v*_*B*_ = *v*_*f*_ − *v*_*r*_, 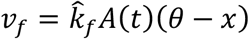 and *v*_*r*_ = *k*_*r*_*x*. *v*_*B*_, *v*_*f*_, and *v*_*r*_ can be seen as a generalized velocity 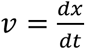.

Equation 4 (part of the replication term in equation 2) describes how daughter cells inherit bound target molecules from the mother cell during replication:

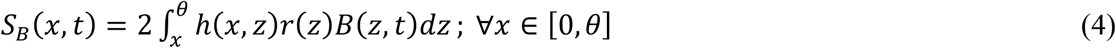

Equation 5 (part of the replication term in equation 2) is a logistic growth model describing reduced bacterial replication as the carrying capacity is approached:

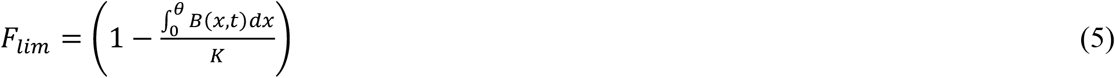

### Model fit to ciprofloxacin time-kill data

We used the quinolone ciprofloxacin to quantitatively fit bacterial time-kill curves, since this is a commonly used antibiotic for which binding parameters have been directly measured. Supplementary Tab. S1 gives an overview of the known parameters used for fitting; Supplementary Tab. S2 gives the parameters resulting from our fit.

The functional relationship between the levels of bacterial replication and death on the fraction of bound target molecules is extremely hard to obtain experimentally. We therefore treated the relationships between the fraction of bound target and bacterial replication and death as free parameters in our model fitting. Ciprofloxacin is considered to have both bacteriostatic and bactericidal action (mixed action)[27, 28], and we fitted functions for a monotonically decreasing replication and a monotonically increasing killing with each successively bound target molecule (see Methods & Supplementary Fig. S1).

Overall, we found that COMBAT could fit the time-kill curves well (R^2^ = 0.93, Fig. 3a). Fig. 3b shows the predicted bacterial replication *r*(*x*) and death as a function of target occupancy *δ*(*x*) based on the fit obtained in Fig. 3a. After model calibration, we simulated bacterial replication during exposure to different antibiotic concentrations for 18 h. For this simulation, positive values indicate an increase in the number of bacteria, and negative values indicate a decrease in the number of bacteria. We estimated a MIC of 0.0139 mg/L (Fig. 3c), a value that is within the range of MIC determinations for wt *E. coli* (0.01 mg/L, 0.015 mg/L, 0.017 mg/L and 0.023 mg/L [11, 29–31]).

**Fig. 3.**
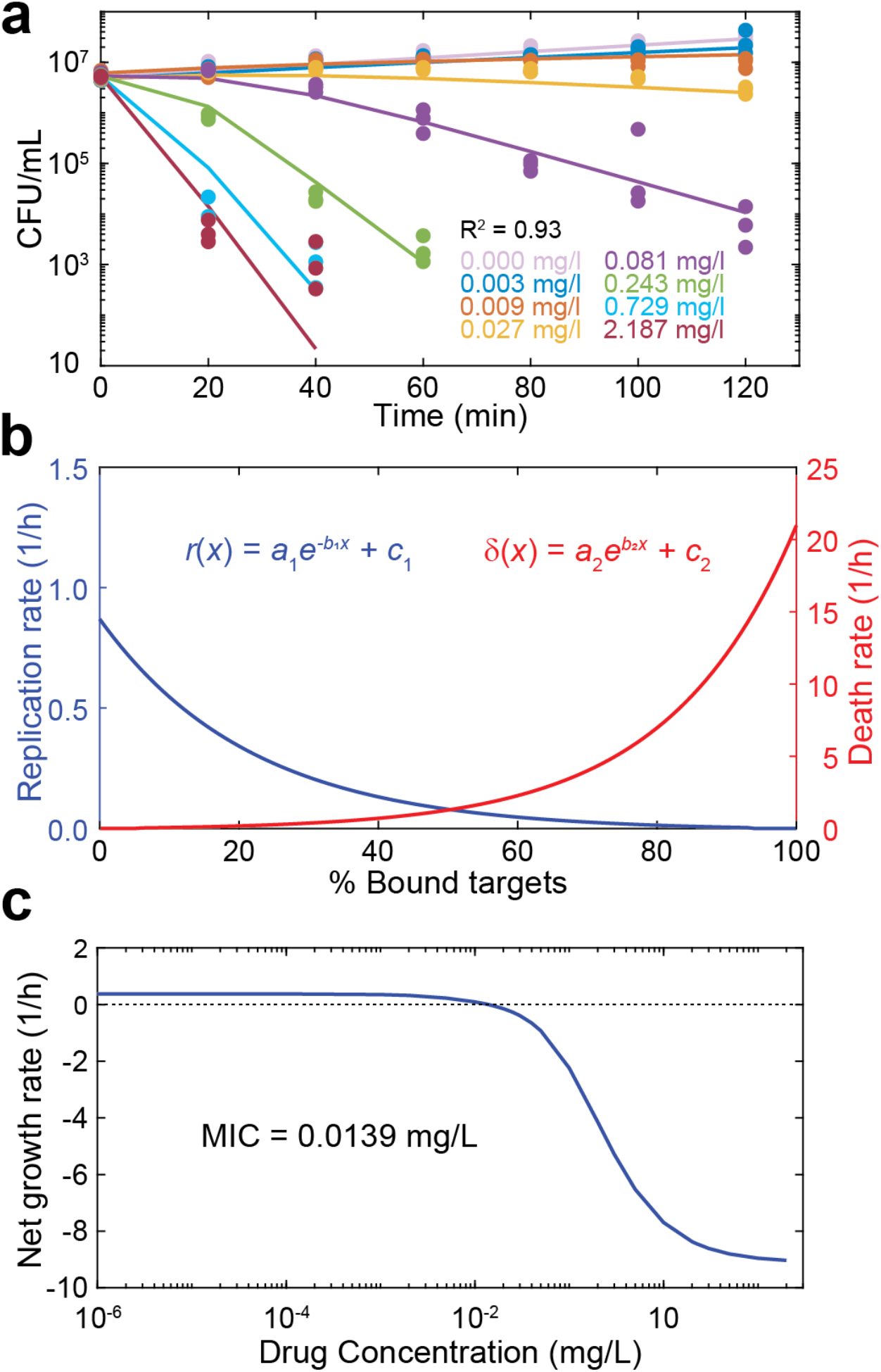
Model predictions for the MIC and the bacteriostatic and bactericidal effects of ciprofloxacin. **a**, Model fit to experimental time-kill curves. The points indicate the experimental data of three independent replicates, and the lines indicate the model fit. Each color indicates a ciprofloxacin concentration as reported in the figure. **b**, The blue line indicates the bacteriostatic effect (*r*(*x*), replication rate) of ciprofloxacin and the red line the bactericidal effect (*δ*(*x*), death rate) as a function of the number of bound targets predicted by the model fit in (**a**). The values of the fitted parameters are listed in Supplementary Tab. S2. **c**, The net growth rate as determined by the slope of a line connecting the initial bacterial density and the final bacterial density of a time-kill curve at 18 h on a logarithmic scale, is given as function of the drug concentration (blue). The dotted horizontal line indicates zero net growth, and the intersection with the blue line predicts the MIC (0.0139 mg/mL).

### Accurate prediction of target overexpression from time-kill data

Having shown that COMBAT can quantitatively fit experimental data on antibiotic action within biologically plausible parameters, we continued to test the predictive ability of the model. Given our hypothesis that modifications in antibiotic-target interactions lead to predictable changes in bacterial susceptibility, we experimentally induced changes in the antibiotic-target interaction of ciprofloxacin in *E. coli*. We then quantified these biochemical changes by fitting COMBAT to corresponding time-kill curves and compared them to the experimental results. Ciprofloxacin acts on gyrase A2B2 tetramers[19]. We used an *E. coli* strain for which both gyrase A and gyrase B are under the control of a single inducible promoter (P_*lacZ*_), such that the amount of gyrase A2B2 tetramer can be experimentally manipulated[32]. We measured net growth rates for this strain at different ciprofloxacin concentrations in the presence of 10 μM isopropyl β-D-1-thiogalactopyranoside (IPTG; mild overexpression) and 100 μM IPTG (strong overexpression) and compared it to the wild-type in the absence of the inducer (Fig. 4a).

**Fig. 4.**
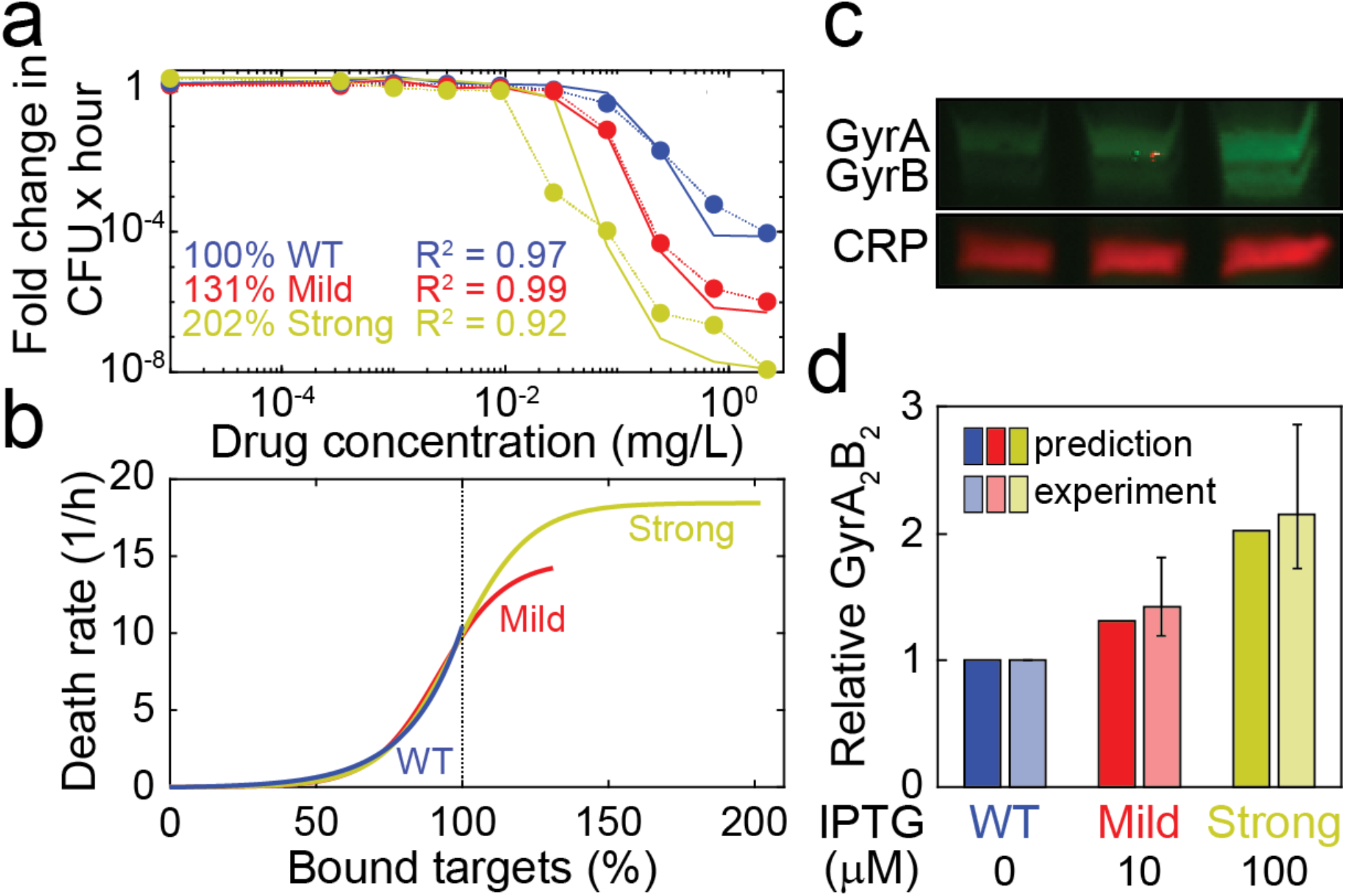
Prediction of relative antibiotic target molecule content from time-kill curves. **a**, Dose-response curves of *E. coli* expressing *gyrA* and *gyrB* under the same IPTG-inducible promoter (SoA3329) grown in the presence of 10 μM IPTG (mild overexpression; red) and 100 μM IPTG (strong overexpression; yellow). A control strain (SoA3330), which expresses wild-type GyrAB levels and contains a mock plasmid, is grown in the absence of inducer (blue). The x-axis indicates the ciprofloxacin concentration, and the y-axis indicates the fold change in colony forming units over time. The dotted lines indicate experimental data, and the solid lines indicate the model fit. The best model fit was obtained for relative target molecule contents of 131 % (mild overexpression) and 202 % (strong overexpression) relative to the control strain (WT). **b**, Death rates of *E. coli* expressing different levels of GyrAB. The colors represent GyrAB expression conditions as in (**a**). The x-axis shows the percentage of bound antibiotic target normalized to the control strain; the y-axis shows the death rate δ(x). Each line represents the best fit for δ(x). **c**, Western blot analysis of GyrA&B in the strains/conditions shown in (**a**). CRP (cAMP receptor protein) was used as loading control. A representative example of six replicates is shown; see Supplementary Fig. S2 for full blots. **d**, comparison of theoretical prediction (from (**b**), solid colors) and GyrA2B2 tetramer levels estimated from relative GyrA&B monomer levels (quantified in (**c**), translucent colors). For the experimental measurements, the bars indicate the mean, and the whiskers represent the 95 % confidence interval.

Like previously reported, we find that increasing gyrase content makes *E. coli* more susceptible to ciprofloxacin[32]. We fitted net growth rates allowing the target molecule content, i.e. gyrase A_2_B_2_, to vary. We assumed that the only change between the different conditions was the amount of target. We further assumed that the relationship between bound target and bacterial replication or death did not differ between the control strain containing a mock plasmid (no IPTG) and the experiments with overexpression (Fig. 4b, between 0 % and 100 %). Finally, we assumed that the maximal kill rate at very high antibiotic concentrations was accurately measured in our experiments and forced the function describing bacterial death through the measured value when all target molecules are bound. We found the best fit for a 1.31x increase in GyrA2B2 target molecule content for bacteria grown in the presence of 10 μM IPTG and a 2.02x increase in GyrA_2_B_2_ target molecule content for those grown in the presence of 100 μM IPTG.

We subsequently tested these predictions experimentally by analyzing Gyrase A and B content by western blot Fig. 4c; Supplementary Fig. S2). Using realistic association and dissociation rates for biological complexes[33], we predicted a range of functional tetramers based on the relative amount of Gyrase A and B proteins (Fig. 4d). Supplementary Tab. S3 details the individual measurements, and the procedure to estimate tetramers is provided in the methods section. We found that the observed overexpression was very close to our theoretical prediction, with 1.43x [95 % CI 1.19-1.81] overexpression (model prediction = 1.31x overexpression) in the presence of 10 μM IPTG and 2.15x [95 % CI 1.73-2.87] overexpression in the presence of 100 μM IPTG (model prediction = 2.02x overexpression).

### Accurate prediction of target occupancy at MIC from time-kill data

Next, we tested whether COMBAT can be applied to the action of the beta-lactam ampicillin, a very different antibiotic with a distinct mode of action from quinolones. Using published pharmacodynamic data of *E. coli* exposed to ampicillin[31] also allowed us to compare COMBAT predictions to established pharmacodynamic approaches. Most of the biochemical parameters for ampicillin binding to its target, penicillin-binding proteins (PBPs), have been determined experimentally (Supplementary Tab. S1). Ampicillin is believed to act as a bactericidal drug[34], and this mode of action is supported by findings from single-cell microscopy[26]. We therefore assume that ampicillin binding does not affect bacterial replication. In order to model the consumption of beta-lactams at target inhibition and eventual target recovery, we made small adjustments to equation 13 (see Methods, description of beta-lactam action).

We fitted COMBAT to published time-kill curves of *E. coli* exposed to ampicillin (Fig. 5a). Again, COMBAT provides a good fit to the experimental data between 0 min and 40-60 min. After that time, observed bacterial killing showed a characteristic slowdown at high ampicillin concentrations which is often attributed to persistence[18] (Fig. 5a). For the sake of simplicity, we chose to omit bacterial population heterogeneity in this work and therefore cannot describe persistence, even though COMBAT can be adapted to capture this phenomenon[18]. Because ampicillin acts in an entirely bactericidal manner, we assume a constant replication rate (see Methods & Supplementary Fig. S1) and fitted bacterial death as a function of target binding, *δ*(*x*) (Fig. 5b, fitted parameters in Tab. S4). Fig. 5c shows the predicted net growth rate over a range of drug concentrations. We estimated a MIC of 2.6 mg/L. This MIC is based on the Clinical & Laboratory Standards Institute definition of the MIC determined at 18 h. The original source of the MIC, which was based on experimental data and a pharmacodynamic model[31] determined an MIC of 3.4 mg/L at 1 h. If we change our prediction to 1 h, our estimated MIC is 3.32 mg/L, which is within 2.5 % of the reported value[31].

**Fig. 5.**
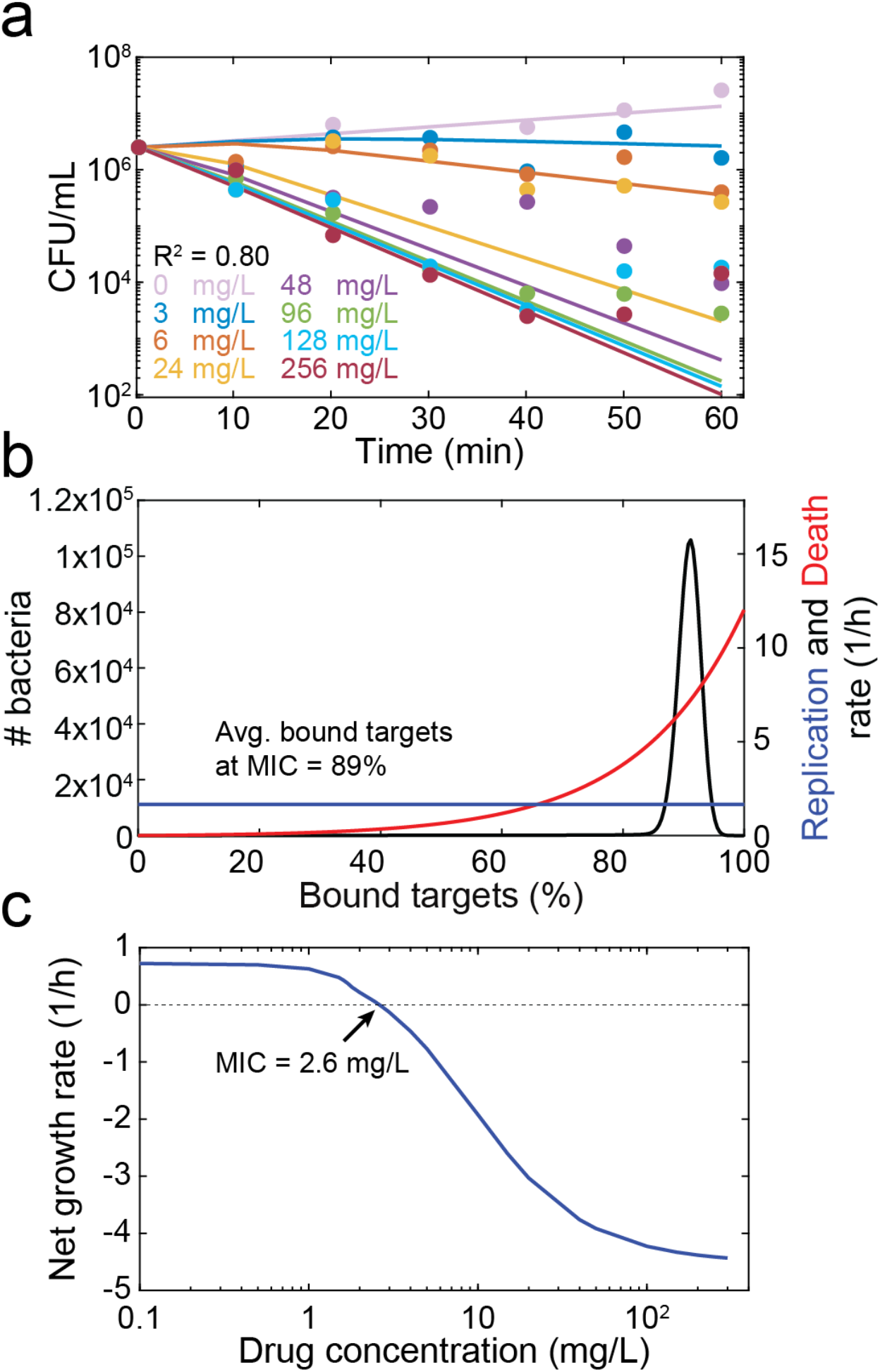
Model prediction of MIC and target occupancy at MIC for ampicillin. **a**, Model fit to previously published time-kill curves[31]. The points represent experimental data, and the lines represent the fit of the model. Each color indicates a single ampicillin concentration, as described in the legend. **b**, Replication (blue) and death (red) rates as a function of the number of bound targets predicted by the model fit in (**a**). The black line indicates the predicted distribution of target occupancies in a bacterial population (both living and dead cells) exposed to ampicillin at the MIC for 18 h. **c**, The net growth rate, as determined by the slope of a line connecting the initial bacterial density and the bacterial density at 18 h on a logarithmic scale predicted from the model fit in (**a**), is shown as function of the drug concentration (blue). The dotted horizontal line indicates zero net growth, and the intersection with the blue line predicts the MIC (2.6 mg/mL).

Having established that COMBAT can also adequately capture the pharmacodynamics of ampicillin, we next tested whether we can estimate experimentally determined target occupancy at the MIC. Our estimated mean occupancy considering both living and dead bacteria is 89 % (Fig. 5b), a value within previously reported experimental estimates from *Staphylococcus aureus* (84-99 %)[35].

### Sensitivity of antibiotic efficacy to parameters of drug-target binding

It is possible to vary all parameters in COMBAT and explore their effect. We used this to test how hypothetical chemical changes to ampicillin or ciprofloxacin would affect antibiotic efficacy (Supplementary Fig. S3-S11). These changes could reflect either bacterial resistance mutations or modifications of the antibiotics themselves. We predict that changes in drug-target affinity, *K*_*D*_, have more profound effects than changes in target molecule content, bacterial reaction to increasingly bound target (i.e. *δ*(*x*) and *r*(*x*)), or changes in target molecule content. We also predict that the individual binding rates *k*_*r*_ and *k*_*f*_, and not just the ratio of these terms, the *K*_*D*_, are important factors in efficiency. The faster a drug binds, the more efficient we predicted it will be. One intuitive explanation for the observation that *k*_*f*_ drives efficacy is that a slow binding fails to rapidly interfere with bacterial replication, which may allow for the production of additional target molecules and thereby reduce the ratio of free antibiotic to target molecules.

### Forecasting the resistance selection window

Finally, we illustrate how COMBAT can be used to explore how the molecular mechanisms of resistance mutations affect antibiotic concentrations at which resistance can emerge, i.e., the resistance selection window. We compared predicted net growth rates as a function of ciprofloxacin concentrations for a wild-type strain and an archetypal resistant strain. For this analysis, we assumed that the resistant strain has a 100x slower drug-target binding rate (i.e. ~100x increased MIC, realistic for novel point mutations[36]) and that the maximum replication rate of the resistant strain is 85 % of the wild type strain[37]. We then predicted the antibiotic concentrations at which resistance would be selected. Interestingly, when comparing COMBAT to previous pharmacodynamics models (Fig. 5), we observed that estimates of replication rates depend on the selected time frame (Fig. 6a). When the timeframe for MIC determination is set to 18 h as defined by CLSI[38], the “competitive resistance selection window”, i.e., the concentration range below the MIC of both strains where the resistant strain is fitter than the wild type, ranges from 0.002 mg/L to 0.014 mg/L for ciprofloxacin (Fig. 6a) and 1 mg/L to 2.6 mg/L for ampicillin (Supplementary Fig. S12), respectively. This corresponds well with previous observations that ciprofloxacin resistance is selected for well below MIC[11]. However, when measuring after 15 min or 45 min, the results are substantially different. The reason for this is illustrated in Fig. 6b. COMBAT reproduces non-linear time kill curves where bacterial replication continues until sufficient target is bound to result in a negative net growth rate. This compares well with experimental data around MIC in Fig. 3a and 5a. In Fig. 6b, we show model predictions for ciprofloxacin concentrations corresponding to a zero net growth (i.e. same population size) after 15 min, 45 min and 18 h (MIC_Resistant; 15 min_, MIC_Resistant; 45 min_, MIC_Resistant; 18 h_). In all cases, the bacterial population first increases and then decreases slowly. This may have consequences for the selection of resistant strains. Fig. 6c illustrates how the resistance selection windows depending on the observed time frame. This suggests that even at concentrations above the 18 h MIC of the resistant strain, there may be initial growth of the resistant strain. In this case, the resistant strain could continue growing at concentration of up to 7 mg/L ciprofloxacin at 15 min, even though the MIC at 18 h is 1.27 mg/L.

**Fig. 6.**
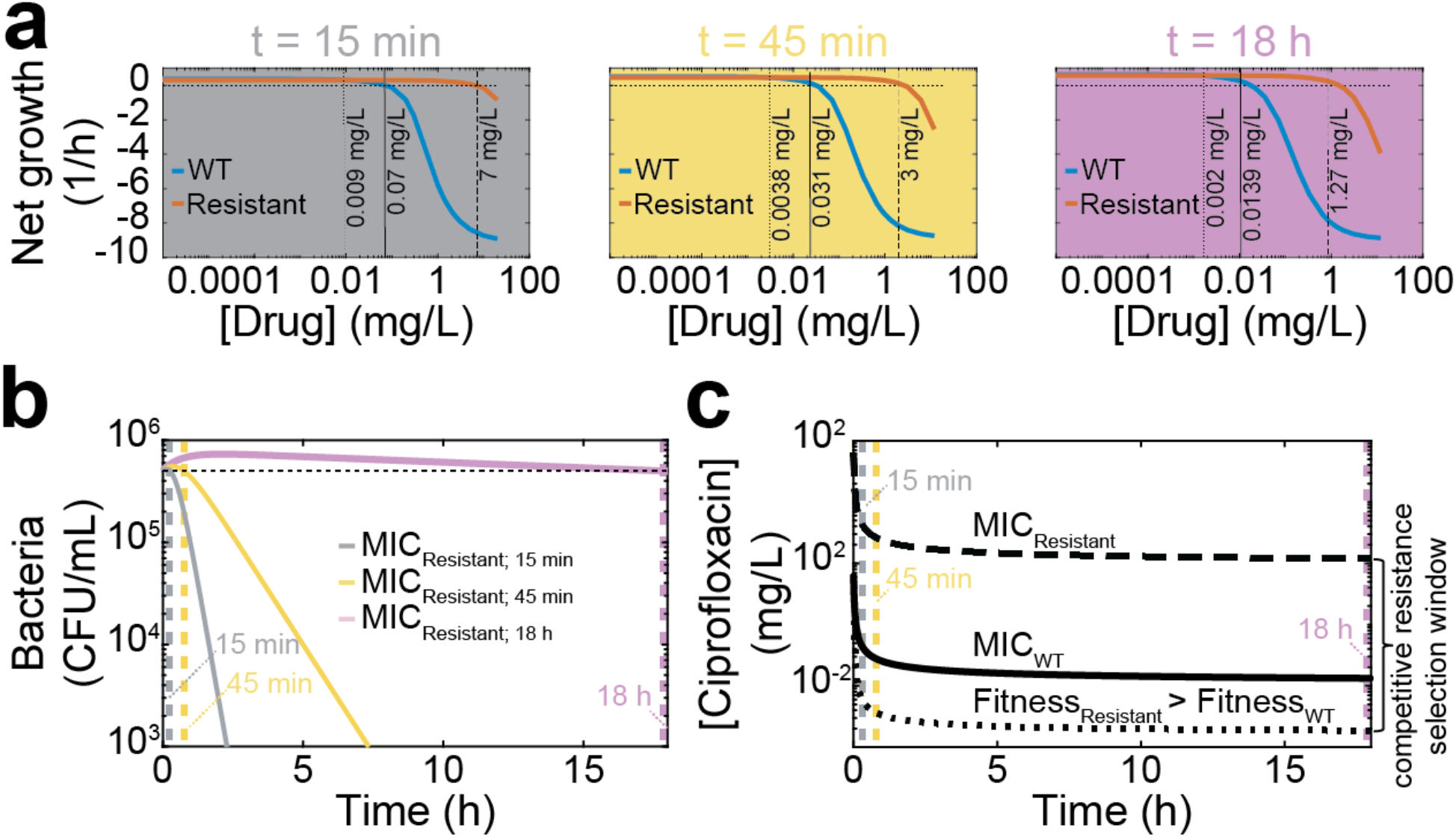
Predicted mutation selection windows for *E. coli* exposed to ciprofloxacin. **a**, The drug concentration of ciprofloxacin is shown on the x-axes, and the average bacterial net growth rate in the first 15 min (grey panel), 45 min (yellow panel), and 18 h (purple panel) of exposure is given on the y-axes. The blue line represents the wild-type strain based on the fits shown in Fig. 3, and the red line represents a strain with a hypothetical resistance mutation that decreases the binding rate (*k*_*f*_) 100-fold and imparts a 15 % fitness cost. The horizontal dotted line indicates no net growth. The vertical dotted line indicates where the resistant strain becomes more fit than the wild-type, the solid vertical line indicates the MIC of the wild-type, and the dashed vertical line indicates the MIC of the resistant strain. **b**, Modeled time kill curves of the resistant strain for ciprofloxacin concentrations at which there is no growth at 15 min (grey; MIC_15 min_ = 7 mg/L), 45 min (yellow; MIC_45 min_ = 3 mg/L) and 18 h (purple; MIC_18 h_ = 1.27 mg/L). The horizontal dotted line indicates the initial population size; the vertical dotted lines represent the time points at which the initial and final population size is the same. **c**, The mutation selection window depends on the time at which bacterial growth is observed. The x-axis shows the observed time at which replication rates were determined, the y-axis shows ciprofloxacin concentrations. The dotted curve shows the ciprofloxacin concentration at which the resistant becomes fitter than the WT (Fitness_Resistant_ > Fitness_WT_), the solid line the MIC of the WT (MIC_WT_), and the dashed line the MIC of the resistant strain (MIC_Resistant_). The area between the dotted and dashed line indicates the competitive resistance selection window.

## Discussion

Optimizing dosing levels of antibiotics is important for maximizing drug efficacy against wild-type strains as well as for minimizing the rise of resistant mutants. The determination of optimal dosing strategies typically requires expensive empirical studies; the need for such studies arises in part from our currently limited capacity to predict how antibiotics will affect bacteria at a given concentration. In fact, drug attrition is mainly due to insufficient predictions of efficacy (pharmacodynamics) rather than pharmacokinetics[6]. For optimizing drug development and for minimizing resistance, we need quantitative predictions for antibiotics or resistant bacterial strains that do not exist yet. The ability to accurately predict MICs on the basis of biochemical parameters and, more generally, to define antibacterial activity across a range of drug concentrations, would allow us to estimate antibiotic efficacy for novel compounds or against not yet emerged resistant strains[15, 39]. Recent studies have reported methods to predict MICs from whole genome sequencing data[40, 41]. However, these methods require transfer of prior knowledge on how the resistance mutations affect MICs in other organisms. There are no methods that could predict *a priori* how chemical changes to an antibiotic structure or novel resistance mutations affect bacterial growth at a given antibiotic concentration.

Here, we accurately predict antibiotic action on the basis of accessible biochemical parameters of drug-target interaction. Our computational model, COMBAT provides a framework to predict the efficacy of compounds based on drug-target affinity, target number, and target occupancy. These parameters may change both when improving antibiotic lead structures as well as when bacteria evolve resistance. Importantly, they can be measured early in drug development and may even be a by-product of target-based drug discovery[42]. When these data are available, COMBAT makes only one assumption: that the rate of bacterial replication decreases and/or the rate of killing increases with successive target binding. While fitting, we allow this relationship to be gradual or abrupt and select the best fit. This means we do not model specific molecular mechanisms down-stream of drug-target binding, but their effects are subsumed in the functions that connect the kinetic of drug-target binding to bacterial replication and death.

In previous work, for example on antipsychotics[16], antivirals[17] and antibiotics[15, 18], models of drug-target binding kinetics have been used to improve our qualitative understanding of pharmacodynamics. Our study substantially advances this work by making accurate quantitative predictions across antibiotics and bacterial strains when measurable biochemical characteristics change. This is possible because COMBAT employs an elegant mathematical approach, based on partial differential equations, that makes it computationally feasible to fit the model to a large range of data. Importantly, we are not only able to predict antibiotic action from biochemical parameters, but can also vice versa use COMBAT to accurately predict biochemical changes from observed patterns of antibiotic action. We have confirmed the excellent predictive power of COMBAT with clinical data as well as experiments with antibiotics with very different mechanisms of action. This gives us confidence that biochemical parameters are major determinants of antibiotic action in bacteria and that COMBAT helps to make rational decisions about antibiotic dosing.

In drug development, our mechanistic modeling approach provides insight into which chemical characteristics of drugs may be useful targets for modification. For example, our sensitivity analyses indicate that antibiotics with a similar affinity but faster binding inactivate bacteria more quickly and therefore prevent replication and production of more target molecules, which would change the ratio of antibiotic to target. Furthermore, because e.g. antibiotic binding and unbinding rates can be determined early in the drug development process, such insight can help the transition to preclinical and clinical dosing trials. This may contribute to reducing bottlenecks between these phases of drug development and thereby save money and time.

Avoiding antibiotic concentrations that select for resistance is challenging because the precise concentrations are only known after extensive experiments have been performed that identify the MIC of (nearly) all possibly emerging resistant mutants. Predicting the resistance selection windows of novel resistant mutants on the basis of biologically plausible changes in drug-target binding would allow us to better assess what drug concentrations need to be achieved to avoid selection of resistance. This approach offers new promise to assess resistance risks prior to characterizing the majority of resistance mutations and thereby reduce the failure rates of candidate compounds late in the drug development process when resistance is observed in patients and substantial resources have been invested.

Our approach also offers insight into determinants of the resistance selection window. Rather than determining the resistance selection window for a comprehensive collection of possibly arising resistance mutations in each bacteria-drug pair, it would be attractive to build transferrable knowledge that allows estimating the resistance selection window. In concordance with a recent meta-analysis of experimental data[43], our sensitivity analyses predict that changes in drug target binding and unbinding have a greater impact on the MIC than changes in target molecule content or down-stream processes. Thus, a more comprehensive characterization of the binding parameters of spontaneous resistant mutants would allow an overview of the maximal biologically plausible levels of resistance that can arise with one mutation. Dosing above this level should then safeguard against resistance. This is especially useful for compounds for which it is difficult to saturate the mutational target for resistance, or for safeguarding against resistance to newly introduced antibiotics for which we do not yet have a good overview of resistance conferring mutations. If toxicity, solubility or other constraints do not allow dosing above the MIC of expected resistant strains, COMBAT can predict the concentration range at which resistance is less strongly selected. This could guide decisions on treating with low versus high doses, which is currently controversially debated[7, 8]. Good quantitative estimates on the dose-response relationship of new drugs would also help defining the therapeutic window, i.e. the range of drug concentrations at which the drug is effective but not yet toxic.

Our quantitative work can help to identify optimal dosing strategies at constant antibiotic concentrations for homogeneous bacterial populations. These measures are commonly used to assess antibiotic efficacy. In addition, previous work has demonstrated that drug-target binding models can qualitatively describe antibiotic efficacy over the fluctuating concentrations that actually occur in patients[26, 44]. They can also explain complicated phenomena such as biphasic kill curves, the post-antibiotic effect, or the inoculum effect[15, 18, 45] that often complicate the clinical phase of drug development. COMBAT has similar characteristics that allow capturing these complex phenomena. Therefore, employing COMBAT may be useful for guiding drug development to maximize antibiotic efficacy and minimize *de novo* resistance evolution.

## Methods

### Mathematical model

COMBAT incorporates the binding and unbinding of antibiotics to their targets and describes how target binding affects bacterial replication and death. This work extends the model developed in[18]. COMBAT consists of a system of two mass balance equations: one PDE for bacteria (describing replication and death as a function of both time and target binding) and one ODE for antibiotic molecules (describing the concentrations as function of time).

In the most basic version of COMBAT, we ignored differences between extracellular and intracellular antibiotic concentrations and only followed the total antibiotic concentration *A*, assuming that the time needed for drug molecules to enter bacterial cells is negligible. We model ciprofloxacin (to which there is a limited diffusion barrier[46]) and ampicillin (where the target is not in the cytosol, even though the external membrane in gram negatives has to be crossed to reach PBPs). We therefore believe that this assumption is justified in wild-type *E. coli*. This basic version of COMBAT is therefore more accurate for describing antibiotic action where the diffusion barrier to the target is weak.

#### Binding kinetics

We describe the action of antibiotics as a binding and unbinding process to bacterial target molecules[18]. For simplicity, we assume a constant number of available target molecules *θ*. The binding process is defined by the formula *A* + *T* ⇌ *x*, where the intracellular antibiotic molecules *A* react with target molecules T at a rate *k*_*f*_ and form an antibiotic-target molecule complex x, where values for x range between 0 and *θ*. If the reaction is reversible, the complex dissociates with a rate *k*_*r*_.

In[18], the association and dissociation terms are described by the following terms

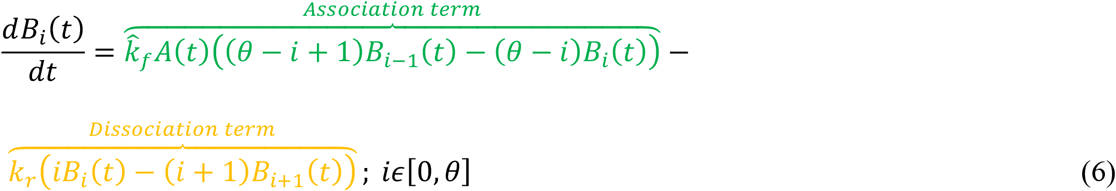

where 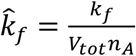, *k*_*f*_ is the association rate, *V*_*tot*_ is the volume in which the experiment is performed, *n*_*A*_ is Avogadro’s number, *k*_*r*_ is the dissociation rate, *B*_*i*_ is the number of bacteria with *i* bound targets, and *θ* is the total number of targets. Green denotes the association term, while the dissociation term is in orange.

This approach requires the use of a large number of ordinary differential equations, (*θ* + 1) for the bacterial population and one for the antibiotic concentration. To generalize this approach, we assume that the variable of bound targets is a real number 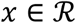. Under this continuity assumption, we consider the bacterial cells as a function of *x* and the time *t*, thereby reducing the total number of equations to two.

Under the continuity approximation 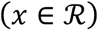, we can rewrite the binding kinetics in the form

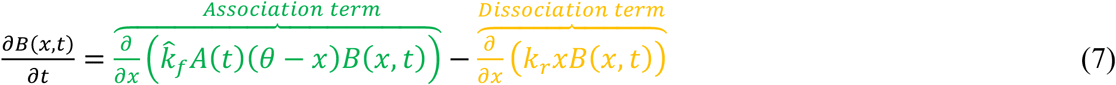

or simply

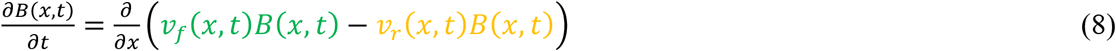

where 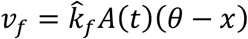 and *v*_*r*_ = *k*_*r*_*x* can be considered as two velocities, i.e., the derivative of the bound targets with respect to the time 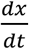. Green denotes the association term, while the dissociation term is in orange.

#### Replication rate

We assume that the replication rate of bacteria, *r*(*x*), is dependent on the number of bound target molecules *x*. The function *r*(*x*) is a monotonically decreasing function of *x*, such that fewer bacteria replicate as more target is bound. *r*(0) is the maximum replication rate, corresponding to the replication rate of bacteria in absence of antibiotics. Thus, *r*(*x*) describes the bacteriostatic action of the antibiotics, i.e., the effect of the antibiotic on bacterial replication.

#### Carrying capacity

Replication ceases as the total bacterial population approaches the carrying capacity *K*. At that point, the replication term of the equation is

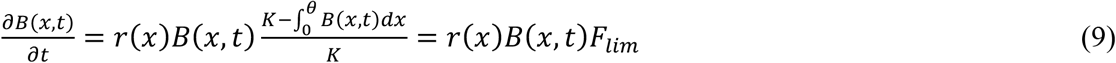

where 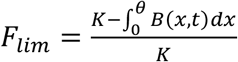 is the replication-limiting term due to the carrying capacity *K*, and 0 ≤ *F*_*lim*_ ≤ 1.

#### Distribution of target molecules upon division

We assume that the total number of target molecules doubles at replication, such that each daughter cell has the same number as the mother cell. We also assume that the total number of drug-target complexes is preserved in the replication and that the distribution of *x* bound target molecules of the mother cell to its progeny is described by a hypergeometric sampling of *n* molecules from *x* bound and 2*θ* − *x* unbound molecules. Under the continuity assumption, we generalize the concept of hypergeometric distribution. Because the hypergeometric distribution is a function of combinations and because a combination is defined as function of factorials, we can use *Γ* functions in place of factorials and redefine a continuous hypergeometric distribution as a function of *Γ* functions. A *Γ* function is

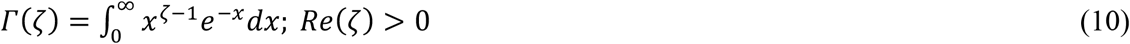

where *ζ* is a complex number. In this way, the distribution can be expressed as a probability density function of continuous variables. The amount of newborn bacteria is given by the term *r*(*x*)*B*(*x*, *t*)*F*_*lim*_(*t*). We assume that bound target molecules are distributed randomly between mother and daughter cells, with each of them inheriting 50% upon division on average. This means that twice the amount of newborn cells must be redistributed along *x* to account for the random distribution process. For example, if a mother cell with 4 bound targets divides, we have two daughter cells, each with a number of bound targets between 0 and 4 (their sum has to be 4), following the generalized hypergeometric distribution. For simplicity, we define *S*(*x, t*) to be a function related to the replication rate that depends on the number of bacteria with a number of bound target molecules ranging between *x* and *θ*, their specific replication rate *r*(*x*), and the fraction of their daughter cells expected to inherit *x* antibiotic-target complexes *h*(*x, z*):

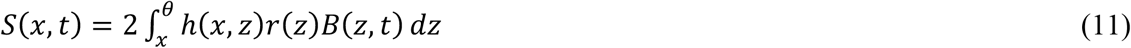

#### Death rate

The death rate function *δ*(*x*) depends on the number of bound target molecules. The function *δ*(*x*) is assumed to be a monotonically increasing function of *x*, where *δ*(*θ*) is the maximum death rate, when all targets in the bacteria have been bound by antibiotics. The shape of this function describes the bactericidal action of the antibiotic.

#### Bacteriostatic and bactericidal effects

We consider several potential functional forms of the relationship between the percentage of bound targets and replication and death rates, because the exact mechanisms how target occupancy affects bacteria is unknown (Supplementary Fig. S1). We use a sigmoidal function that can cover cases ranging from a linear relationship to a step function. When the inflection point of a sigmoidal function is at 0 % or 100 % target occupancy, the relationship can also be described by an exponential function. We assume that replication in bactericidal and death in bacteriostatic drugs is independent of the amount of bound target. With sufficient experimental data, the replication rate *r*(*x*) and/or the death rate *δ*(*x*) can be obtained by fitting COMBAT to time-kill curves of bacterial populations after antibiotic exposure. The sigmoidal shape of *r*(*x*) and *δ*(*x*) can be written as:

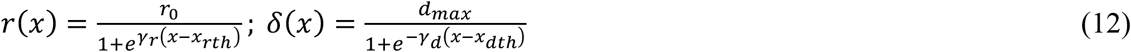

where *x*_*rth*_ is the replication rate threshold, *x*_*dth*_ is the death rate threshold, and both represent the point where the sigmoidal function reaches ½ of its maximum. γ_r_ and γ_d_ represent the shape parameters of the replication and death rate functions, respectively. These factors determine the steepness around the inflection point. When they are extreme, the relationship approaches a linear or a step function.

#### Full equation describing bacterial population

Putting these components together, the full equation describing a bacterial population is:

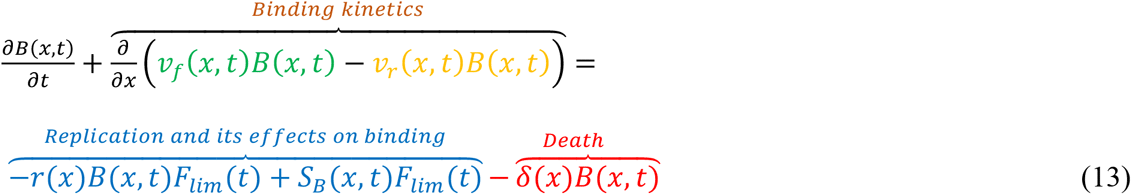

where *B*(*x, t*) is the number of bacteria. As in equations 2, 6, 7 and 8, green denotes the binding term, orange the unbinding term (together the binding kinetics is given in brown), blue the term describing bacterial replication and red the term describing bacterial death.

#### Equation describing antibiotic concentration

The free antibiotic concentration results from mass conservation, i.e., all antibiotic molecules associating with their target are subtracted and all dissociating antibiotic molecules are added. Equation 3 in the results section describes the dynamics of the antibiotic concentration.

#### Description of beta-lactam action

Beta-lactams acetylate their target molecules (PBPs) and thereby inhibit cell wall synthesis. The acetylation of PBPs consumes beta-lactams. However, PBPs can recover through deacetylation. We modified the term of drug-target dissociation in the equation describing antibiotic concentrations (equation 3), and set the unbinding rate *k*_*r*_ = 0. To reflect the recovery of target molecules, we substituted the dissociation rate *k*_*r*_ in the equation describing the bacterial population with the deacetylation rate *k*_*a*_, as described in[26].

#### Initial and boundary conditions

At *t* = 0, we assume that all bacteria have zero bound targets (*x* = 0), and the initial concentration of bacteria is *B*(*x*, 0) = 0, *x* > 0, and *B*(0,0) = *B*_0_.

At the boundaries of the partial differential equation (*x* = 0, *x* = *θ*), we specify that the outgoing velocities are zero. For *x* = 0, i.e. no bound target molecules, the unbinding velocity *v*_*r*_(0, *t*) = 0, and in *x* = *θ*, i.e. all targets are bound, the binding velocity *v*_*f*_(*θ*, *t*) = 0. When the replication term at *x* = 0 and the death term at *x* = *θ* are known, we can solve the partial differential equation with two ordinary differential equations at the boundaries. They are similar to the equations at *x* = 0 and at *x* = *θ* described by Abel zur Wiesch et al.[18], but taking into account that x is a continuous variable instead of a natural number.

#### Numerical schemes

To solve our system of differential equations, we used a first-order upwind scheme. Specifically, we used the spatial approximation 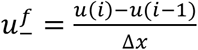 for the binding term (*v*_*f*_ > 0) and the spatial approximation 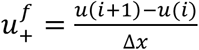 for the unbinding term (*v*_*r*_ < 0). For the time approximation of both the PDEs and the ODEs, we used the forward approximation 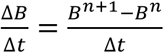[47]. We also verified that the Courant-Friedrichs-Lewy condition is satisfied. For fitting the experimental data of bacteria exposed to ciprofloxacin and ampicillin, we used the particle swarm method (“particleswarm” function in Matlab, MathWorks software).

#### Concentrations of gyrase A_2_B_2_ tetramers

We assumed that gyrases A and B first homo-dimerize to A_2_ and B_2_, respectively, which in turn bind to each other to form the tetramer TR[48]. The following system of equations describes their binding kinetics:

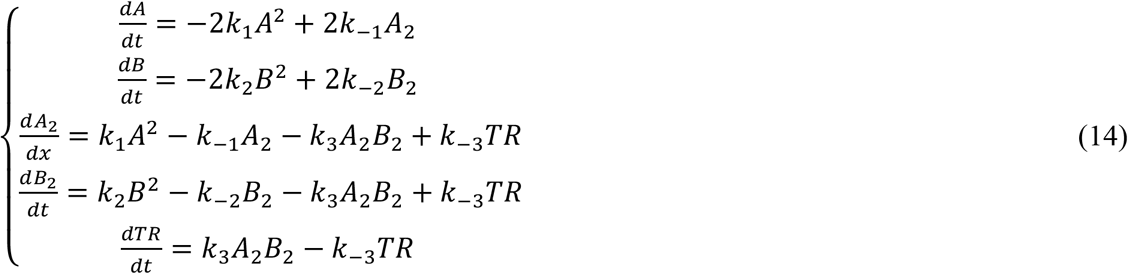

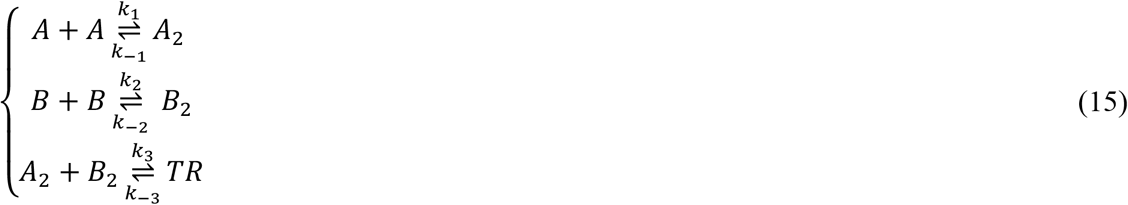

First, we calibrated the model to ensure that we obtain the correct number of gyrase A_2_B_2_ tetramers (~100) per wild type bacterial cell[49, 50]. This results in an average of each 206 gyrase A and B monomers. Because the association and dissociation rates of the dimers and tetramers are unknown, we sampled 10^4^ sets of six parameters in equation 14 (*k*_−3_,…*k*_3_) in a Latin hypercube approach from a biologically plausible range where the association rates are between 10^7^ - 10^9^ M^−1^ s^−1^ and the dissociation rates between 10^−3^ - 10^−1^ s^−1^ [33]. This results in 10^4^ estimates for each of the six experimental replicates quantifying gyrase A and B (Fig. 4, Supplementary Fig. S2, Supplementary Tab. S3).

### Experimental methods

#### Strains, growth conditions and strain construction

*Escherichia coli* strain BW25113[51] (SoA2740) was transformed with plasmids pCA24N-SC101-gyrAB[32] and pCA24N-SC101-ΔP-YFP[32] using electroporation, resulting in strains BW25113/pCA24N-SC101-gyrAB (SoA3329) and BW25113/pCA24N-SC101-ΔP-YFP (SoA3330), respectively. pCA24N-SC101-gyrAB encodes the *E. coli gyrAB* genes under control of the IPTG inducible LacZ promoter. pCA24N-SC101-ΔP-YFP encodes a promoterless copy of YFP and was used as a control. Bacteria were grown at 30°C on either LB agar or in LB broth, both supplemented with 10 μg/mL chloramphenicol (Cm) and 10 μM (mild induction) or 100 μM (strong induction) of isopropyl β-D-1-thiogalactopyranoside (IPTG) (43714 5X, VWR Chemicals) when necessary.

#### Time-kill curves

Overnight cultures of BW25113 or SoA3329 and SoA3330 were diluted 1:1000 in pre-warmed LB or LB-Cm and LB-Cm-IPTG, respectively, and grown with shaking to OD_600_ ~0.5. A 1:3 dilution series of ciprofloxacin was made and added to the cultures at indicated concentrations. Additional cultures without antibiotics and with a very high concentration of ciprofloxacin (2.187 mg/L) were used to determine the minimal and maximal kill rate, respectively. Samples were taken immediately prior to addition of the antibiotic and in ~20 min intervals or after 45 min, respectively. Samples were washed once in phosphate buffered saline (PBS) before colony forming units (CFUs) were determined for each sample by plating a 1:10 dilution series in PBS on LB agar plates.

#### GyrAB quantification

To quantify the relative amount of GyrAB, samples of SoA3329 and SoA3330 were collected after 45 min of drug treatment as described above. An equal number of cells corresponding to 1 mL culture at OD_600_ = 1 were harvested by centrifugation. Pelleted cells were lysed at room temperature for 20 min using B-PER bacterial protein extraction reagent (90078, Thermo Scientific) supplemented with 100 μg/mL lysozyme, 5 units/mL DNaseI (all part of B-PER™ with Enzymes Bacterial Protein Extraction Kit, 90078, Thermo Scientifc) and 100 μM/mL PMSF (52332, Calbiochem). Samples were stored at −80°C until further use.

Samples were heated to 70°C for 10 min after addition of 1x Bolt sample reducing agent (B0009, Life Technologies) and 1x fluorescent compatible sample buffer (LC2570, Invitrogen). Proteins in whole-cell lysates were separated on 4-15 % Mini-Protean TGX Precast gels (456-1085, Bio-Rad) and transferred to 0.2 μm Nitrocellulose membranes (1704158, Bio-Rad).

Membranes were blocked in Odyssey blocking buffer-TBS (927-50000, Li-Cor) for at least one hour at room temperature. Primary antibodies raised against GyrA (Rabbit α-Gyrase A, PA005, Inspiralis), GyrB (Rabbit α-Gyrase B, PB005, Inspiralis), and CRP (Mouse α-*E. coli* CRP, 664304, Nordic Biosite antibodies) were diluted 1:250, 1:250, and 1:2,000 in Odyssey blocking buffer-TBS, respectively. The blocked membranes were incubated with the appropriate primary antibodies overnight at 4°C, washed 4x for 15 min each in TBS-T solution (Tris buffered saline supplemented with Tween20: 0.138 M sodium chloride, 0.0027 M potassium chloride, 0.1 % Tween20, pH 8.0 at 25°C), and incubated for 2 h at room temperature with fluorescent labelled secondary antibodies (1:10,000 of IRDye® 680RD Goat anti-Mouse IgG, P/N 925-68070, Li-Cor and 1:5000 of IRDye® 800CW Goat anti-Rabbit IgG, P/N 925-32211, Li-Cor) in Odyssey blocking buffer-TBS. Finally, the membranes were washed 4x for 15 min each in TBS-T solution and imaged at 700 nm and 800 nm using a Li-Cor Odyssey Sa scanning system. Band intensities were quantified from unmodified images using the record measurement tool of Photoshop CS6, normalized to the CRP loading control after background subtraction, and reported relative to SoA3330. For clarity, the “levels” tool of Photoshop CS6 was used to enhance the contrast of shown Western blot images.

## Data Availability

Computer code will be available at https://www.abel-zur-wiesch-lab.com/.

## Acknowledgements

We thank Jingyi Liang, Vi Tran, Antal Martinecz, Giovanni Montani, Forrest W. Crawford, Roberto Natalini, Klaus Harms, Christoph Zimmer, Angelo Vannozzi and Rafal Mostowy for helpful discussions and feedback on the manuscript. We would like to acknowledge R. Regoes for providing previously published raw data. This work was funded by Bill and Melinda Gates Foundation Grant OPP1111658 (to T.C. & P.AzW.), Research Council of Norway (NFR) Grant 262686 (to P.AzW.) and 249979 (to S.A.), and Helse-Nord Grant 14796 (to S.A.).

## Author Contributions

P.AzW. designed the study. F.C. developed the mathematical models. A.P., B.S., M.S., and S.A. designed the experiments. A.P., B.S., M.S., and S.L. performed the experiments. F.C., A.P., B.S., M.S., T.C., S.A., and P.AzW. analyzed the data. T.C., S.A., and P.AzW. wrote the manuscript.

## Competing Interests statement

The authors declare no competing interests.

