## Supplementary_Information for "Drug-target binding quantitatively predicts optimal antibiotic dose levels"

| Antibiotic | Target | Parameter | Value | References |
|---|---|---|---|---|
| Ciprofloxacin | Gyrase | Copy #/cell | Gyrase: 50-100 functional tetramers | [1, 2] |
| | | Unbinding rate $k_r$ | $3\cdot10^{-4}$ sec$^{-1}$ | [3] |
| | | $K_D$ ($k_r$/ $k_f$) | $10^{-5.6}$<br>$10^{-5.7}$<br>$10^{-5.9}$<br>$10^{-7.5}$ *(lower boundary according to authors)* | [4]<br>[5]<br>[6]<br>[3] |
| Ampicillin | Penicillin binding proteins | Copy #/cell | ~ 2500±120 | [7, 8] |
| | | Binding rate $k_f$ | 130 | [9, 10] |
| | | Deacetylation rate $k_a$ | $10^{-4}$ | [10] |

**Supplementary Tab. S1I Kinetic parameters for used antibiotics.** To our knowledge, the association rate for ciprofloxacin has not been determined directly. Because the values of the ratio of dissociation rate $k_r$ and association rate $k_f$, $K_D$, diverge by more than an order of magnitude in the literature, we chose to fit the association rate $k_f$ as a free parameter in our model while constraining $K_D$ to remain within the published range (the resulting value of $k_f$ is given together with other fitted parameters in Supplementary Tab. S2).

| Parameter | Value | Unit of measure | Explanation |
|---|---|---|---|
| $k_f$ | $3.210 \cdot 10^{3}$ | $M^{-1}sec^{-1}$ | Binding rate |
| $r_0$ | $2.416 \cdot 10^{-4}$ | $sec^{-1}$ | Maximum replication WT |
| $a_1$ | $2.441 \cdot 10^{-4}$ | $sec^{-1}$ | Coefficient of r(x) |
| $b_1$ | 0.0459 | - | Coefficient of r(x) |
| $c_1$ | $-2.478 \cdot 10^{-6}$ | $sec^{-1}$ | Coefficient of r(x) |
| $\delta_m$ | 0.0058 | $sec^{-1}$ | Maximum death rate |
| $a_2$ | $2.46 \cdot 10^{-5}$ | $sec^{-1}$ | Coefficient of δ(x) |
| $b_2$ | 0.0547 | - | Coefficient of δ(x) |
| $c_2$ | $-2.46 \cdot 10^{-5}$ | $sec^{-1}$ | Coefficient of δ(x) |

**Supplementary Tab. S2l Parameters resulting from model fit to experimental time-kill curves of ciprofloxacin in wild-type *E. coli*.** Fig. 3a shows the model fit and Fig. 3b shows the resulting functions for the replication rate $r(x)$ and the death rate $\delta(x)$ as a function of the number of bound targets.

| GyrA (m) | GyrB (m) | GyrA$_2$B$_2$ (m) | GyrA (s) | GyrB (s) | GyrA$_2$B$_2$ (s) |
|---|---|---|---|---|---|
| 1.67 | 1.85 | 1.7136 ± 0.0004 | 2.43 | 3.46 | 2.5017 ± 0.001 |
| 1.09 | 1.20 | 1.1121 ± 0.002 | 1.44 | 1.68 | 1.4803 ± 0.0004 |
| 1.11 | 1.21 | 1.1673 ± 0.0002 | 1.56 | 1.94 | 1.6024 ± 0.0005 |
| 2.04 | 2.99 | 2.1004 ± 0.0009 | 3.36 | 6.65 | 3.4706 ± 0.0016 |
| 1.16 | 1.26 | 1.1802 ± 0.0002 | 1.77 | 2.29 | 1.8229 ± 0.0007 |
| 1.25 | 1.51 | 1.2797 ± 0.0004 | 1.98 | 3.04 | 2.0401 ± 0.0008 |
| Mean + IC 95% | | 1.43 (1.119 – 1.81) | Mean + IC 95% | | 2.15 (1.73 – 2.87) |

**Supplementary Tab. S3l Results of gyrase level determination and estimated GyrA$_2$B$_2$ tetramer levels**. (m) indicates mild overexpression, (s) indicates strong overexpression. Columns headed GyrA and GyrB show the experimentally determined overexpression as fold expression compared to the wild type. The columns headed GyrA$_2$B$_2$ show the estimated tetramer levels resulting from each measurement. For the GyrA$_2$B$_2$ tetramer estimation, we sampled $10^4$ sets association and dissociation rates from a uniform distribution within their reported limits (Latin hypercube approach). We report the standard deviation for each estimate. We give summary estimates in the last row of the table.

| Parameter | Value | Unit of measure | Explanation |
|---|---|---|---|
| $r_0$ | $4.664 \cdot 10^{-4}$ | $sec^{-1}$ | Maximum replication WT |
| $a_3$ | $1.075 \cdot 10^{-5}$ | $sec^{-1}$ | Coefficient of δ(x) |
| $b_3$ | 0.023 | - | Coefficient of δ(x) |
| $c_3$ | $-1.075 \cdot 10^{-6}$ | $sec^{-1}$ | Coefficient of δ(x) |
| $\delta_m$ | 0.0034 | $sec^{-1}$ | Maximum death rate |

**Supplementary Tab. S4l Parameters resulting from model fit to experimental time-kill curves of ampicillin in wild-type *E. coli*.** Fig. 5b shows the resulting death rate $\delta(x)$ as an exponential function of the number of bound targets $\delta(x) = a_3 e^{b_3 x} + c_3$.

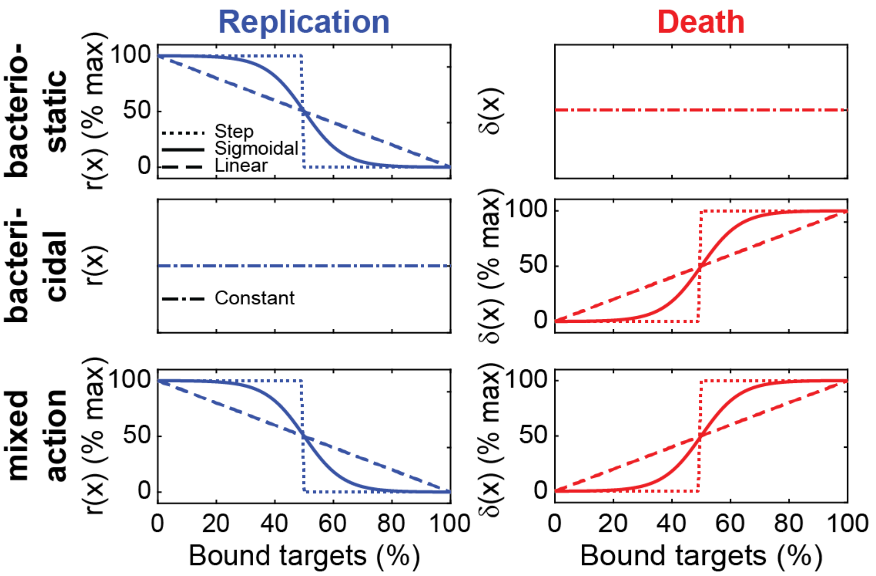


**Supplementary Fig. S1l Functions connecting successive antibiotic target binding with bacterial replication and death in bacteriostatic and bactericidal drugs.** These graphs show which functions were used to fit the dependence of bacterial replication, $r(x)$, and death, $\delta(x)$, on target occupancy for antibiotics with bacteriostatic, bactericidal or mixed action. Solid lines indicate a sigmoidal relationship, dotted lines indicate a step function, dashed lines indicate a linear relationship, and dash-dotted lines indicate independence, i.e. a constant replication or death rate. The left panels, show the replication rates (blue), the right panels show the death rates (red). The top panels, show rates for a bacteriostatic drug, the middle panels, show rates for a bactericidal drug, and the bottom panels, show rates for a drug with mixed effects. The sum of the replication and death rates at certain target occupancies gives the net growth or decline rate of the bacterial population.

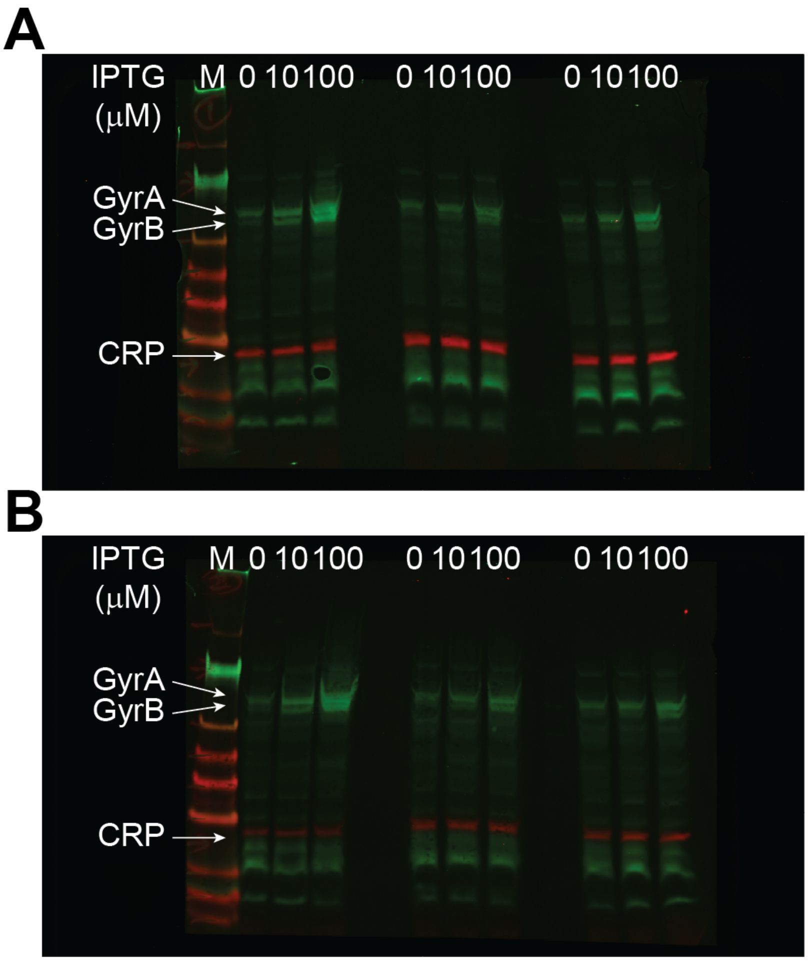


**Supplementary Fig. S2l Western blots of GyrA, GyrB and CRP in *E. coli* whole cell lysates. a, b**, *E. coli* expressing *gyrA* and *gyrB* under control of the same IPTG-inducible promoter (SoA3329) grown in the presence of 10 $\mu$M IPTG (mild overexpression) and 100 $\mu$M IPTG (strong overexpression). A control strain containing a mock plasmid (SoA3330), representing wild-type GyrAB levels, was grown in the absence of inducer. Whole cell lysates were separated on a SDS-PAGE gel, blotted, and detected with specific fluorescent antibodies against GyrA (green), GyrB (green) and CRP (red). CRP (cAMP receptor protein) was used as loading control.

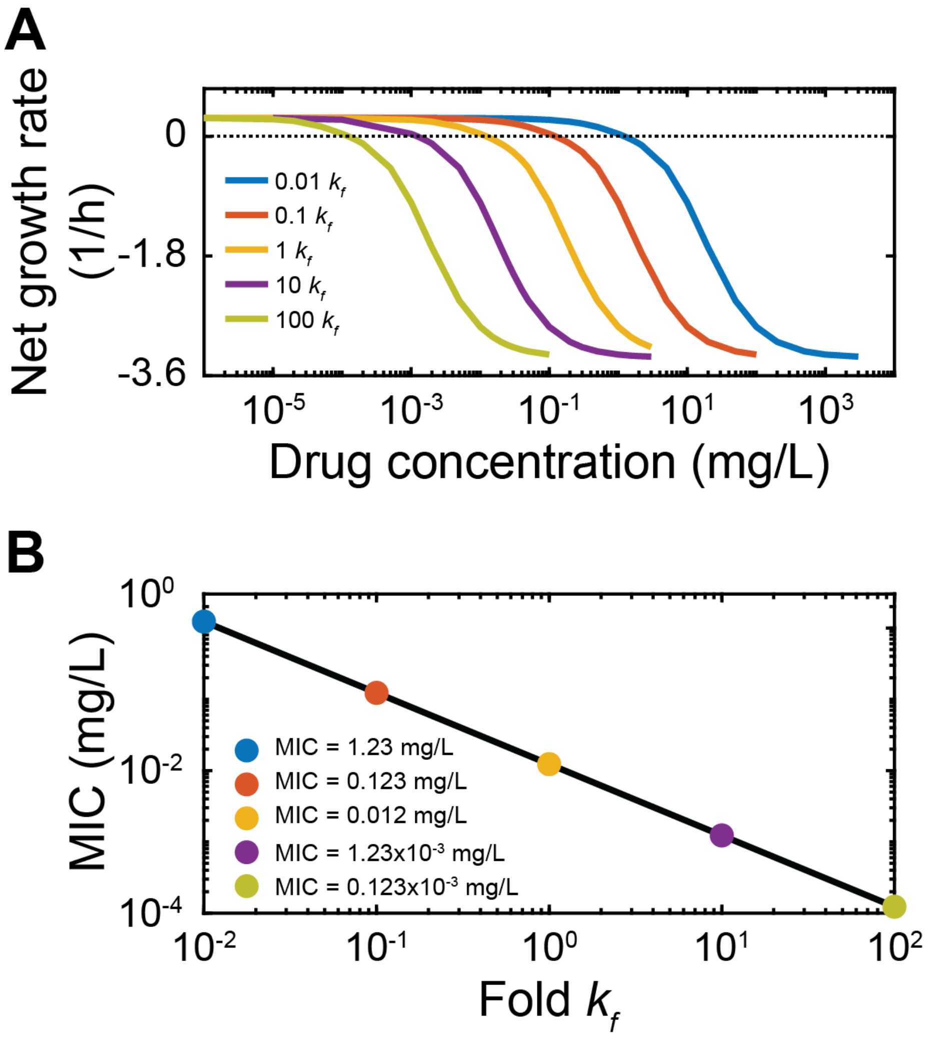


**Supplementary Fig. S3l Sensitivity analysis of ciprofloxacin fit: changes in the binding rate $k_f$.** We use the model fitted to experimental data to explore the sensitivity of our results to changes in $k_f$ (0.01x, 0.1x, 1x, 10x, and 100x original value). **a**, Net growth rate ($\log_{10}$(bacterial number at 18 h) - $\log_{10}$(bacterial number at 0 h))/18 h) as function of drug concentration for different values of the binding rate $k_f$ (see legend). The dotted horizontal line indicates zero net growth. The intersections of the simulated dose-response curves with this line indicate the corresponding MICs. **b**, Sensitivity of the MIC to $k_f$ obtained from simulations in (**a**). The color code indicates the MIC corresponding to the simulation with the same color in (**a**).

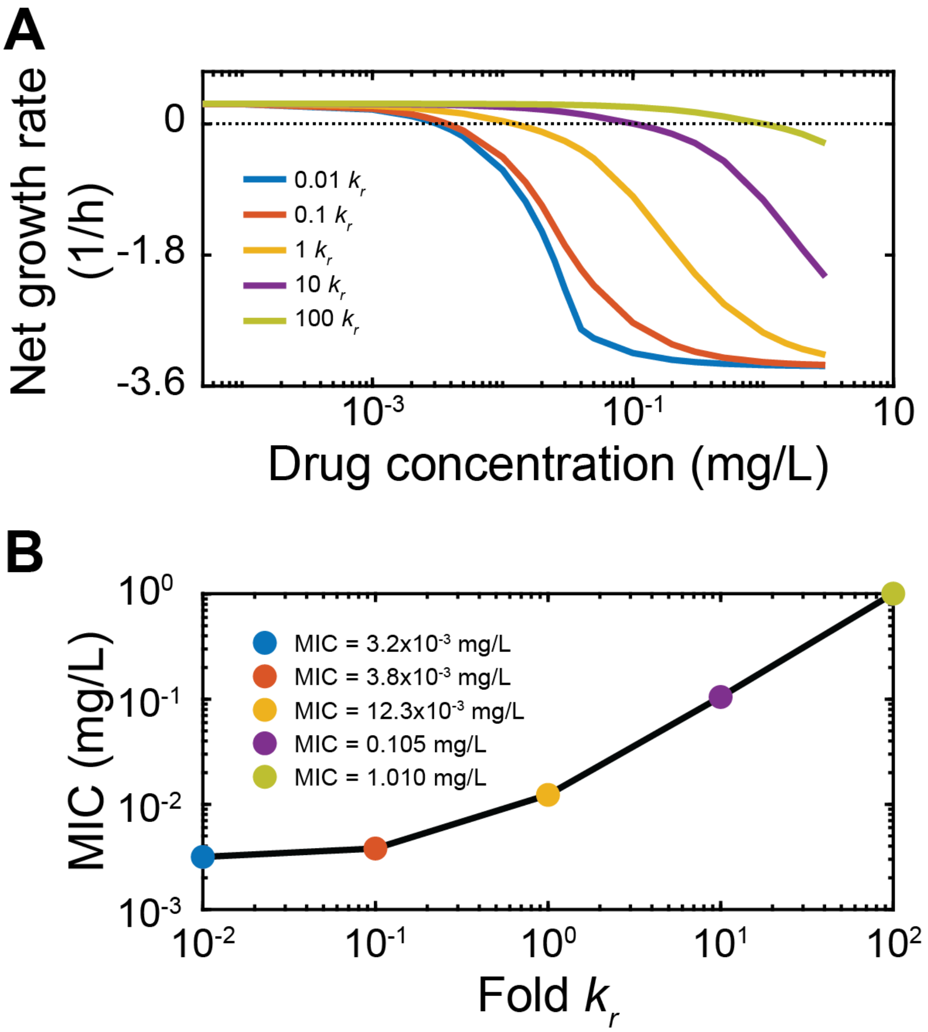


**Supplementary Fig. S4l Sensitivity analysis of ciprofloxacin fit: changes in the dissociation rate $k_r$.** We use the model fitted to experimental data to explore the sensitivity of our results to changes in $k_r$ (0.01x, 0.1x, 1x, 10x, and 100x original value). **a**, Net growth rate ($\log_{10}$(bacterial number at 18 h) - $\log_{10}$(bacterial number at 0 h))/18 h) as function of drug concentration for different values of the binding rate $k_r$ (see legend). The dotted horizontal line indicates zero net growth. The intersections of the simulated dose-response curves with this line indicate the respective MICs. **b**, Sensitivity of the MIC to $k_r$ obtained from simulations in (**a**). The color code indicates the MIC corresponding to the simulation with the same color in (**a**).

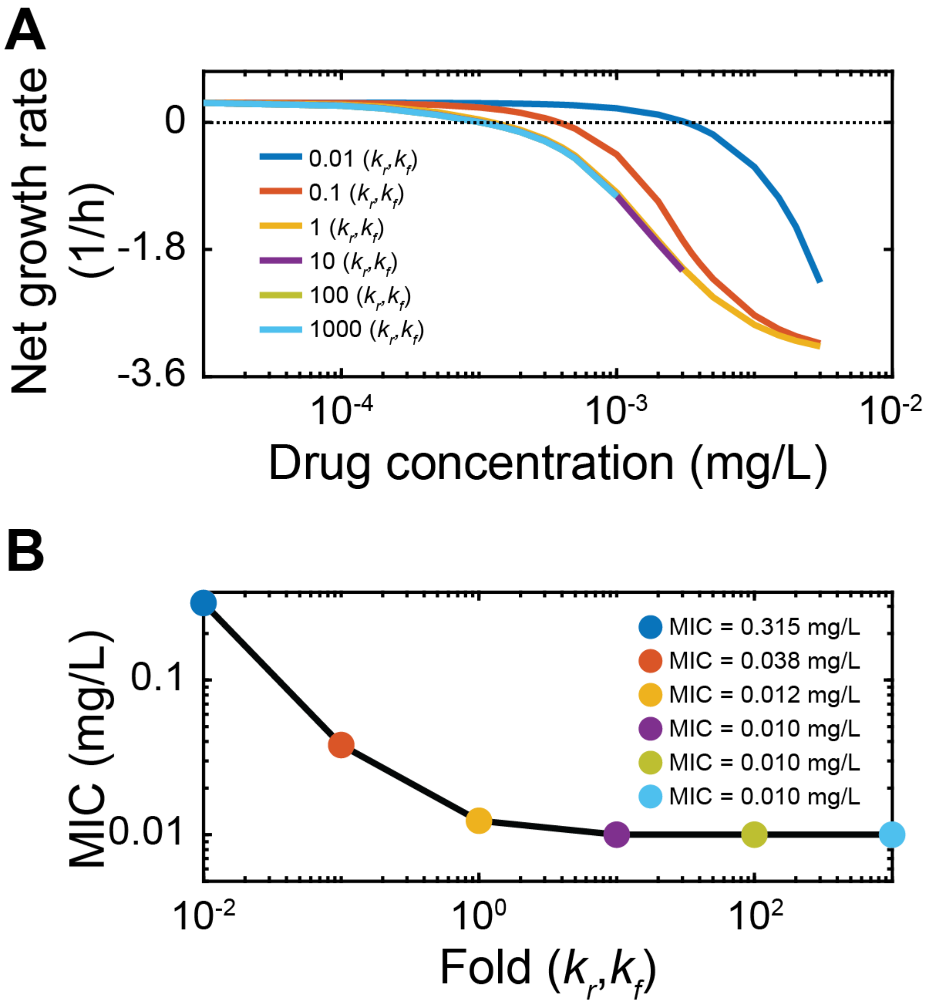


**Supplementary Fig. S5l Sensitivity analysis of ciprofloxacin fit: changes in the drug-target turnover rate.** We use the model fitted to experimental data to explore the sensitivity of our results to changes in the turnover rate of the drug-target complex. We changed $k_r$ and $k_f$ (0.01x, 0.1x, 1x, 10x, 100x, and 1000x original value) while keeping the ratio between $k_f$ and $k_r$, the affinity $K_D$, constant. **a**, shows the net growth rate ($\log_{10}$(bacterial number at 18 h) - $\log_{10}$(bacterial number at 0 h))/18 h) as function of drug concentration for different values of the turnover rate. The dotted horizontal line indicates zero net growth. The intersections of the simulated dose-response curves with this line indicate the respective MICs. **b**, Sensitivity of the MIC to turnover rate obtained from simulations in (**a**). The color code indicates the MIC corresponding to the simulation with the same color in (**a**).

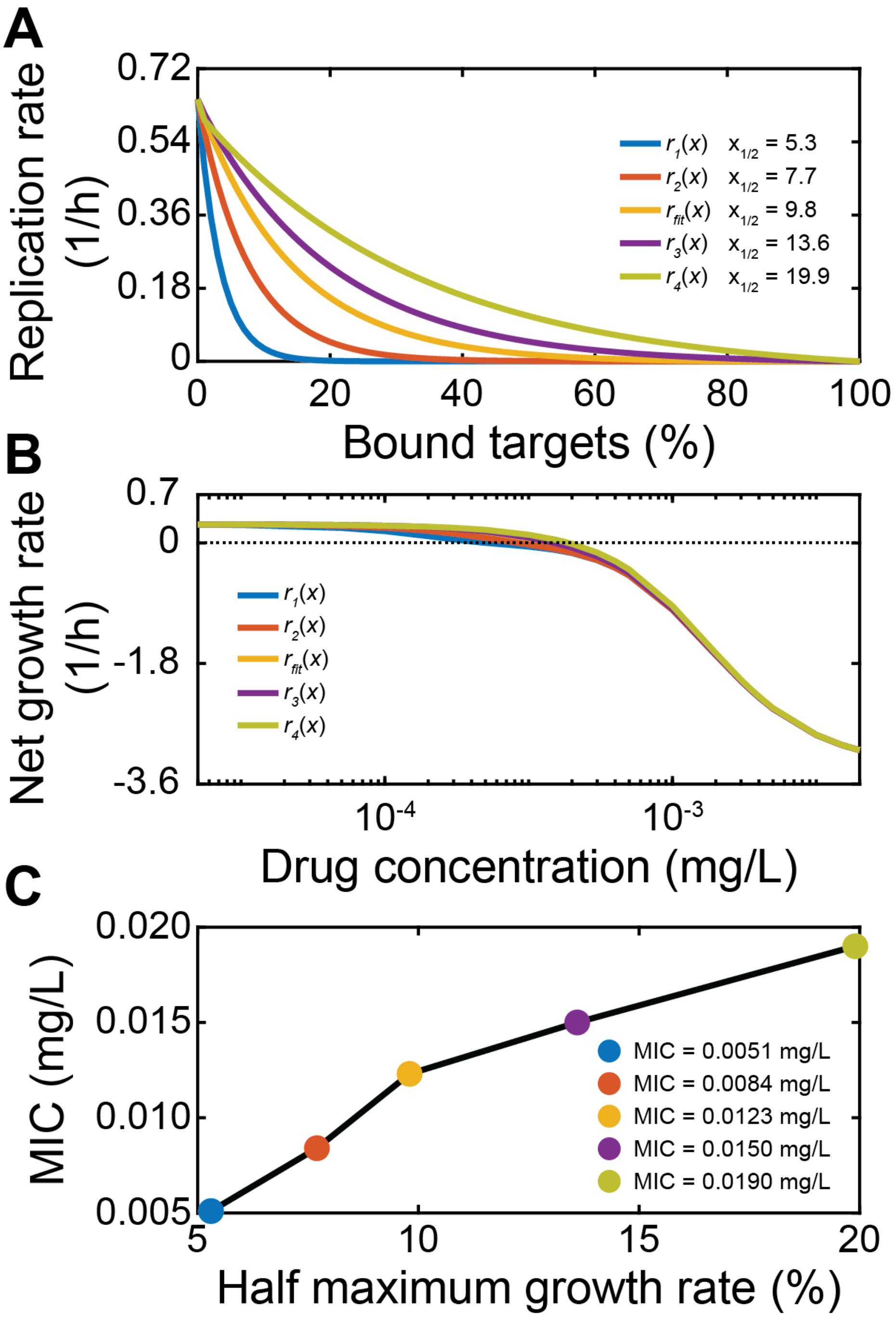


**Supplementary Fig. S6| Sensitivity analysis of ciprofloxacin fit: changes in the replication rate with increasingly bound target *r*(x).** We use the model fitted to experimental data to explore the sensitivity of our results to changes in the replication rate with increasingly bound target *r*(x). We change the value of bound target at which we obtain a half-maximal replication rate, $x_{1/2}$. **a**, Functions connecting bacterial replication rates *r*(x) to percentage of bound target molecules with different half-maximal replication rates. **b**, Net growth rate ($\log_{10}$(bacterial number at 18 h) - $\log_{10}$(bacterial number at 0 h))/18 h) as function of drug concentration for different values of $x_{1/2}$ (see legend). The dotted horizontal line indicates zero net growth. The intersections of the simulated dose-response curves with this line indicate the respective MICs. **c**, Sensitivity of the MIC to $r(x)$ obtained from simulations in (**b**). The color code indicates the MIC corresponding to the simulation with the same color in (**a**&**b**).

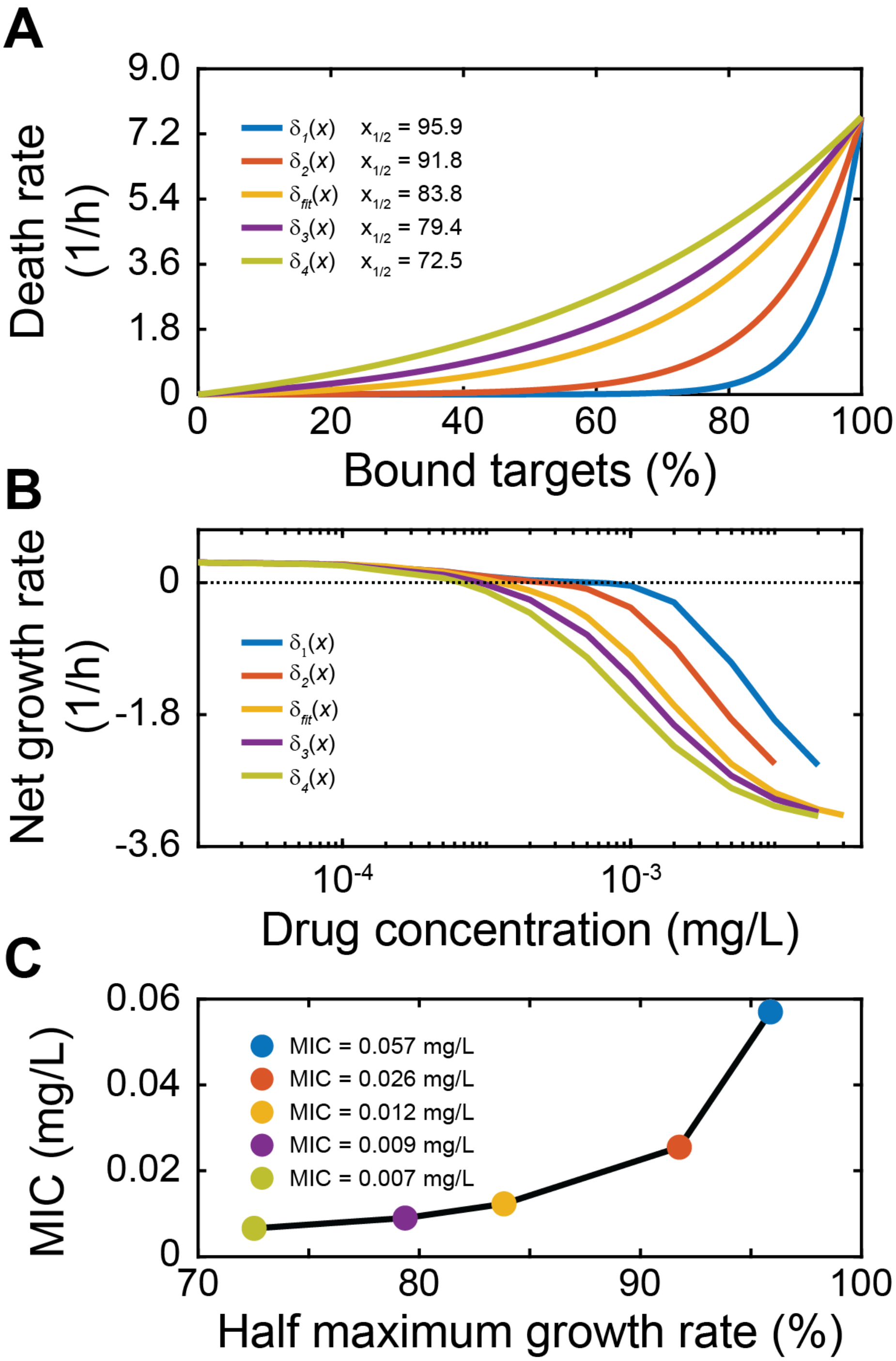


**Supplementary Fig. S7I Sensitivity analysis of ciprofloxacin fit: changes in the death rate with increasingly bound target** $\boldsymbol{\delta(x)}$**.** We use the model fitted to experimental data to explore the sensitivity of our results to changes in the death rate with increasingly bound target $\delta(x)$. We change the value of bound target at which we obtain a half-maximal death rate, $x_{1/2}$. **a**, Functions connecting bacterial death rates $\delta(x)$ to percentage of bound target molecules with different half-maximal death rates. **b**, Net growth rate ($\log_{10}$(bacterial number at 18 h) - $\log_{10}$(bacterial number at 0 h))/18 h) as function of drug concentration for different values of $x_{1/2}$ (see legend). The dotted horizontal line indicates zero net growth. The intersections of the simulated dose-response curves with this line indicate the respective MICs. **c**, Sensitivity of the MIC to $\delta(x)$ obtained from simulations in (**b**). The color code indicates the MIC corresponding to the simulation with the same color in (**a**&**b**).

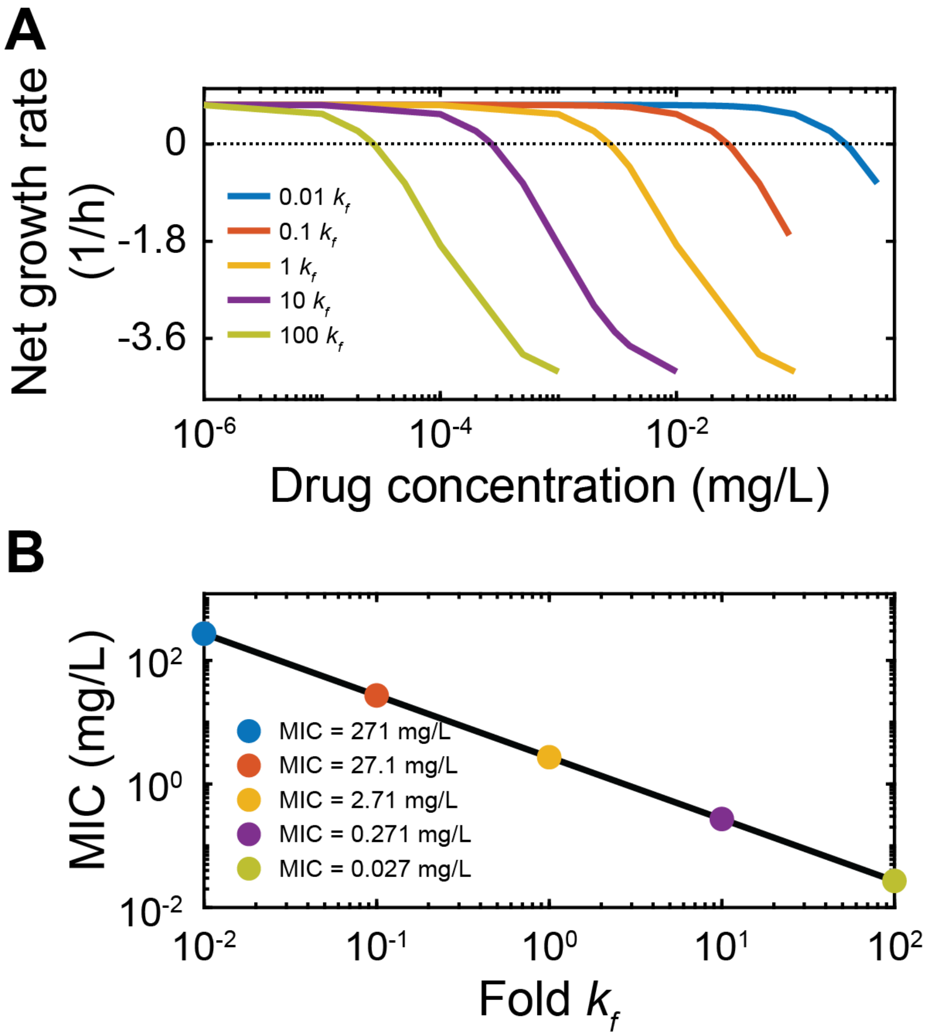


**Supplementary Fig. S8l Sensitivity analysis of ampicillin fit: changes in the binding rate $k_f$.** We use the model fitted to experimental data (Fig. 5) to explore the sensitivity of our results to changes in $k_f$ (0.01x, 0.1x, 1x, 10x, and 100x original value). **a**, Net growth rate ($\log_{10}$(bacterial number at 18 h) - $\log_{10}$(bacterial number at 0 h))/18 h) as function of drug concentration for different values of the binding rate $k_f$ (see legend). The dotted horizontal line indicates zero net growth. The intersections of the simulated dose-response curves with this line indicate the respective MICs. **b**, Sensitivity of the MIC to $k_f$ obtained from simulations in (**a**). The color code indicates the MIC corresponding to the simulation with the same color in (**a**).

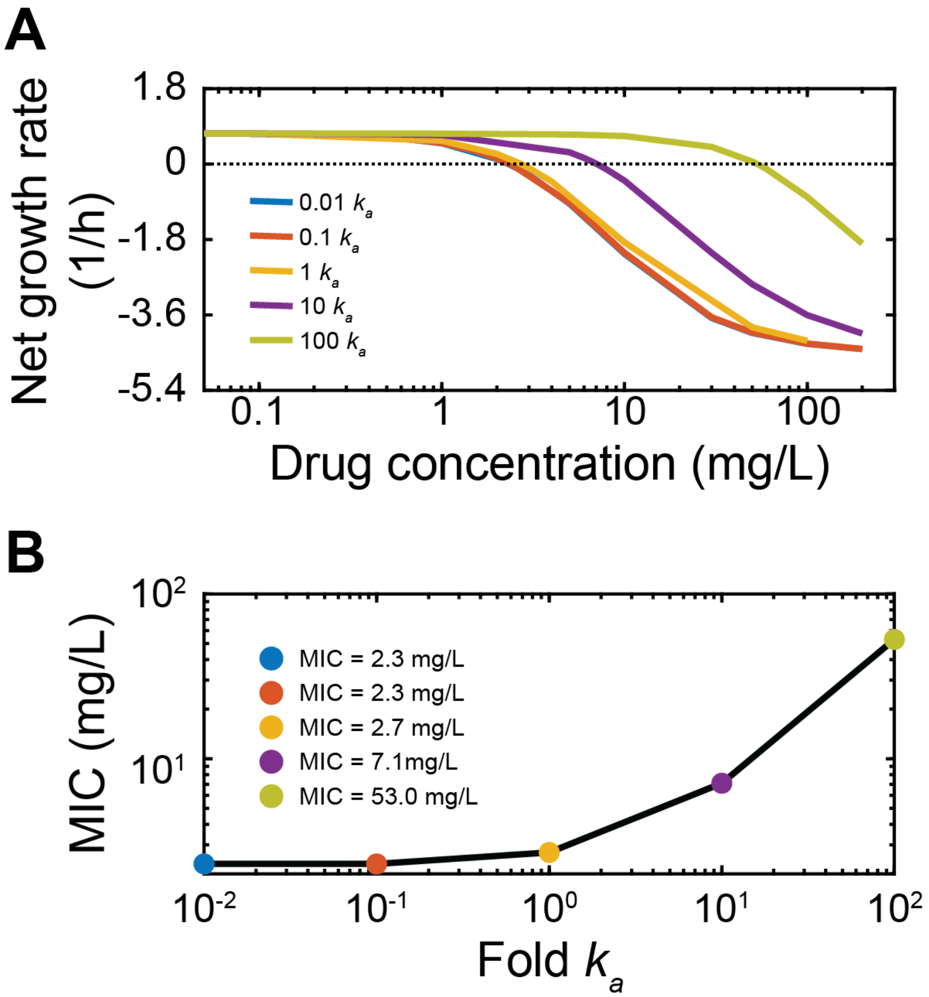


**Supplementary Fig. S9l Sensitivity analysis of ampicillin fit: changes in the deacetylation rate $k_a$.** We use the model fitted to experimental data (Fig. 5) to explore the sensitivity of our results to changes in $k_a$ (0.01x, 0.1x, 1x, 10x, and 100x original value). **a**, Net growth rate ($\log_{10}$(bacterial number at 18 h) - $\log_{10}$(bacterial number at 0 h))/18 h) as function of drug concentration for different values of the binding rate $k_a$ (see legend). The dotted horizontal line indicates zero net growth. The intersections of the simulated dose-response curves with this line indicate the respective MICs. **b**, Sensitivity of the MIC to $k_a$ obtained from simulations in (**a**). The color code indicates the MIC corresponding to the simulation with the same color in (**a**).

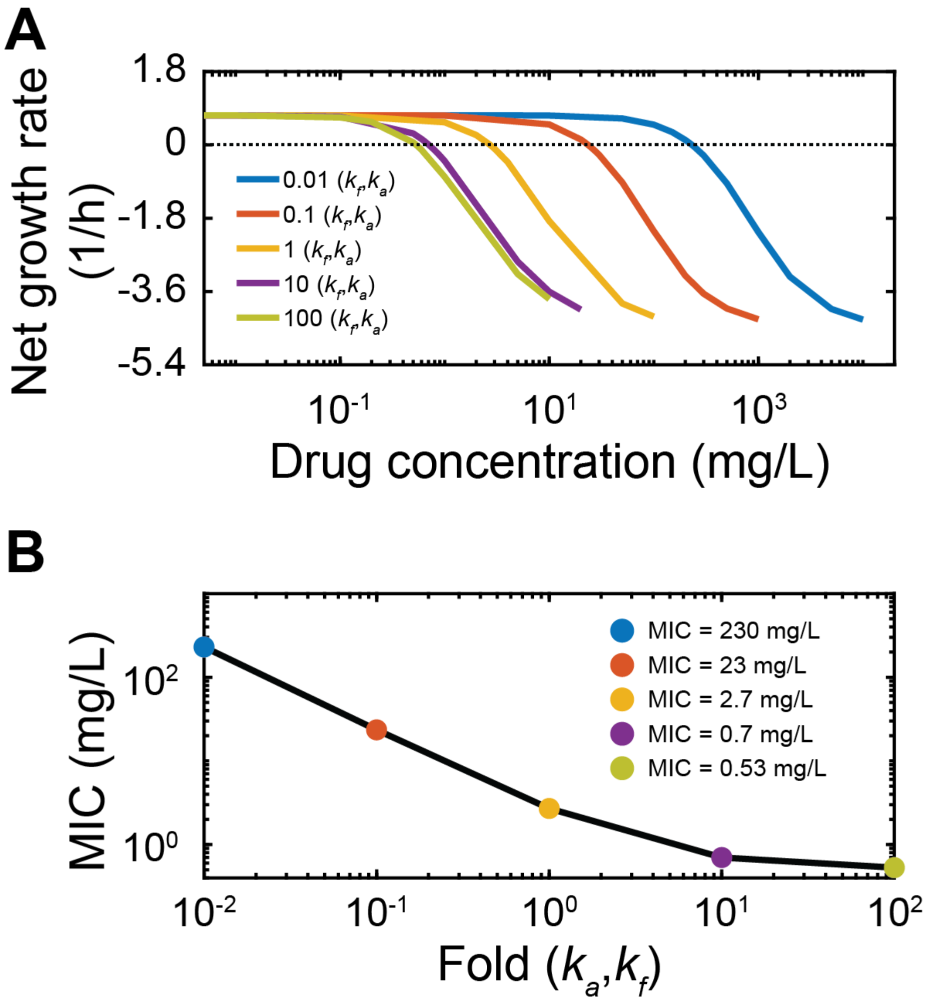


**Supplementary Fig. S10l Sensitivity analysis of ampicillin fit: changes in the drug-target turnover rate.** We use the model fitted to experimental data (Fig. 5) to explore the sensitivity of our results to changes in the turnover rate of the drug-target complex. We changed values for $k_a$ and $k_f$ (0.01x, 0.1x, 1x, 10x, and 100x original value) while keeping the ration of $k_a/k_f$ constant. **a**, Net growth rate ($\log_{10}$(bacterial number at 18 h) - $\log_{10}$(bacterial number at 0 h))/18 h) as function of drug concentration for different values of the turnover rate (see legend). The dotted horizontal line indicates zero net growth. The intersections of the simulated dose-response curves with this line indicate the respective MICs. **b**, Sensitivity of the MIC to turnover rate obtained from simulations in (**a**). The color code indicates the MIC corresponding to the simulation with the same color in (**a**).

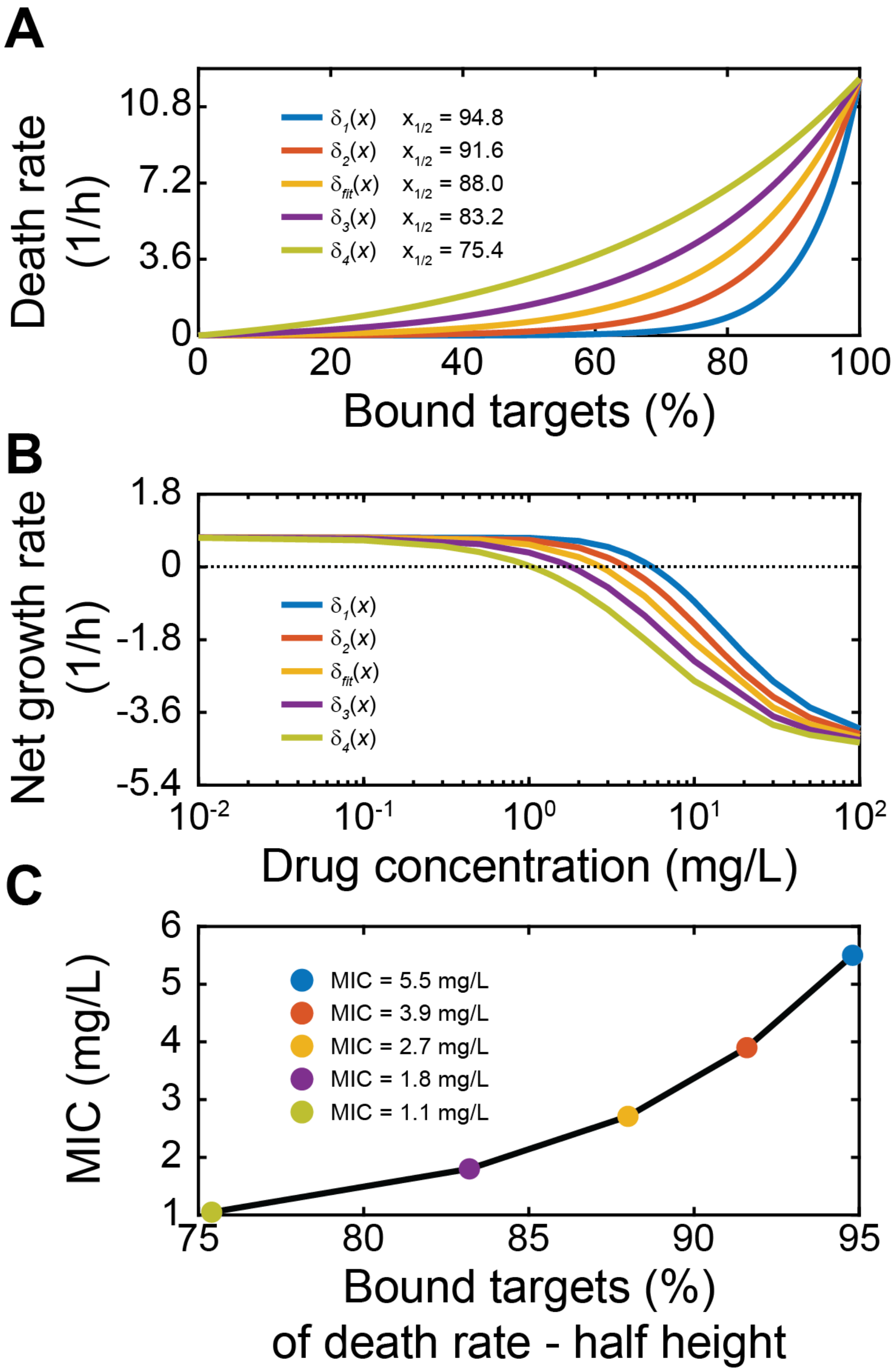


**Supplementary Fig. S11l Sensitivity analysis of ampicillin fit: changes in the death rate with increasingly bound target $\delta(x)$.** We use the model fitted to experimental data (Fig. 5) to explore the sensitivity of our results to changes in the death rate with increasingly bound target $\delta(x)$. We change the value of bound target at which we obtain a half-maximal death rate, $x_{1/2}$ (see legend). **a**, Functions connecting bacterial death rates $\delta(x)$ to percentage of bound target molecules with different half-maximal replication rates $x_{1/2}$ (see legend). **b**, Net growth rate ($\log_{10}$(bacterial number at 18 h) - $\log_{10}$(bacterial number at 0 h))/18 h) as function of drug concentration for different values of $\delta(x)$ (see legend). The dotted horizontal line indicates zero net growth. The intersections of the simulated dose-response curves with this line indicate the respective MICs. **c**, Sensitivity of the MIC to $\delta(x)$ obtained from simulations in (**b**). The color code indicates the MIC corresponding to the simulation with the same color in (**a**&**b**).

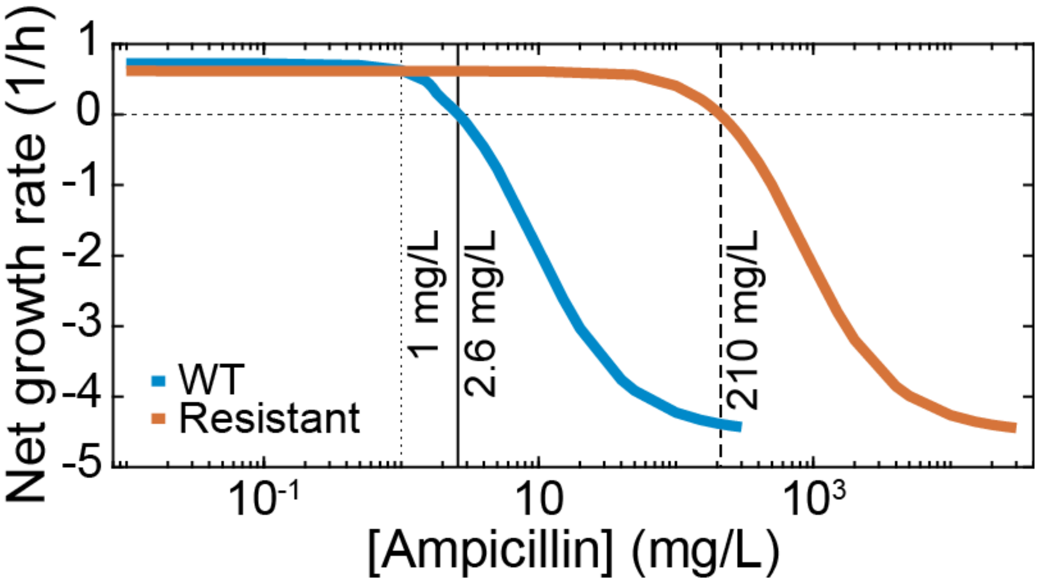


**Supplementary Fig. S12l Predicted mutation selection windows for *E. coli* exposed to ampicillin.** The drug concentration of ampicillin is shown on the x-axes, and the average bacterial net growth rate over 18 h is given on the y-axes. The blue line represents the wild-type strain based on the fits shown in Fig. 5, and the red line represents a strain with a theoretical resistance mutation that decreases the binding rate ($k_f$) 100-fold and imparts a 15% fitness cost. The dotted horizontal line represents no net growth. The first vertical dotted line indicates where the resistant strain becomes fitter than the wild-type (the start of the competitive resistance selection window), the solid vertical line indicates the MIC of the wild-type (the start of the classical resistance selection window), and the dashed vertical line indicates the MIC of the resistant strain, above which selection for resistance should be minimal because both growth of the wild-type and the resistant strain is inhibited.
